# Re-defining how mRNA degradation is coordinated with transcription and translation in bacteria

**DOI:** 10.1101/2024.04.18.588412

**Authors:** Seunghyeon Kim, Yu-Huan Wang, Albur Hassan, Sangjin Kim

**Affiliations:** Department of Physics, University of Illinois at Urbana-Champaign, Urbana, IL 61801, USA; Center for Biophysics and Quantitative Biology, University of Illinois at Urbana–Champaign, Urbana, IL 61801, USA

**Keywords:** mRNA degradation, RNase E, transcription-translation coupling, premature transcription termination, transertion, co-transcriptional regulatcanion

## Abstract

In eukaryotic cells, transcription, translation, and mRNA degradation occur in distinct subcellular regions. How these mRNA processes are organized in bacteria, without employing membrane-bound compartments, remains unclear. Here, we present generalizable principles underlying coordination between these processes in bacteria. In *Escherichia coli*, we found that co-transcriptional degradation is rare for mRNAs except for those encoding inner membrane proteins, due to membrane localization of the main ribonuclease, RNase E. We further found, by varying ribosome binding sequences, that translation affects mRNA stability not because ribosomes protect mRNA from degradation, but because low translation leads to premature transcription termination in the absence of transcription-translation coupling. Extending our analyses to *Bacillus subtilis* and *Caulobacter crescentus*, we established subcellular localization of RNase E (or its homolog) and premature transcription termination in the absence of transcription-translation coupling as key determinants that explain differences in transcriptional and translational coupling to mRNA degradation across genes and species.

## Introduction

Unlike eukaryotic cells, bacterial cells do not have a nucleus, and the transfer of genetic information from DNA to protein takes place within a common space, the cytoplasm, permitting concurrence of translation and even mRNA degradation while mRNA is transcribed^1,2^. How transcription, translation, and mRNA degradation are coordinated during the life cycle of an mRNA, in the absence of membrane-bound compartments, is a fundamental question that underpins gene expression regulation in bacterial cells. Addressing this question will enable understanding of how protein expression levels are regulated by the cell in response to different environments^3–5^ and also inform the design of synthetic gene expression systems, where precise manipulation of gene expression is essential^6^.

Decades of research in a model bacterium, *E. coli*, has provided strong evidence that transcription is coupled to translation^7–11^, that mRNA degradation can start during transcription^12–15^, and that translation affects mRNA degradation^16,17^. However, whether this picture is broadly applicable across all genes and different bacterial species remains unclear. Identifying the molecular and sequence variables affecting coupling between transcription, translation, and mRNA degradation will lead to a generalizable model for understanding the regulation of bacterial gene expression. In this work, we investigated when (during the life cycle of mRNA) and where (within a cell) mRNAs are degraded in coordination with transcription and translation in bacterial cells and identified key factors that contribute to commonalities and differences in co-transcriptional and post-transcriptional control of gene expression in *E. coli*, *B. subtilis*, and *C. crescentus*.

The possibility of co-transcriptional mRNA degradation has been discussed since early studies of long operons in *E. coli*. In *lac* and *trp* operons, mRNA sequences from the promoter-proximal gene were shown to decay before the promoter distal genes were transcribed^12–14^. A genome-wide measurement of mRNA lifetimes in *E. coli* compared transcription elongation time and mRNA lifetime and suggested that many long genes that exhibit transcription elongation times longer than mRNA lifetimes may experience co-transcriptional mRNA degradation^15^. Co-transcriptional mRNA degradation can have a significant impact by reducing the number of proteins made per transcript, which can be beneficial when a quick stop in protein synthesis is needed to respond to changing cellular needs. However, whether co-transcriptional degradation is indeed possible in *E. coli* remains in question because the main ribonuclease controlling mRNA degradation, RNase E, is found on the inner membrane of the cell, away from the nucleoid^18–21^. Co-transcriptional mRNA degradation would therefore need to invoke the dynamic relocalization of a gene locus to the membrane, which has been observed for certain genes^22,23^. Furthermore, unlike in *E. coli*, RNase E is localized in the cytoplasm in *C. crescentus*^24–26^, raising a question about how mRNA degradation is differentially controlled in *C. crescentus* cells in comparison to *E. coli* cells.

In contrast to the lack of clarity how mRNA degradation can be coupled to transcription in bacterial cells, several studies have supported the coupling of mRNA degradation to translation, such that mRNAs with a strong ribosome binding sequence have long lifetimes^16,17,27–31^. This trend has been explained by the notion that ribosomes protect mRNA from degradation. However, what aspect of ribosome activity—for example, whether it is the rate of loading at the 5’ end of the mRNA or whether it is ribosomal density across the mRNA—is responsible for the protective role remains unclear^16,17,32^. Understanding the exact mechanism of translation that affects mRNA lifetime would help make quantitative predictions for expression output for different genes.

In this work, we used *lacZ* as a model gene to study how transcription, translation, and mRNA degradation are coordinated in bacterial cells. The lac operon in *E. coli* is a paradigm of bacterial gene regulation, and our current understanding of transcription-translation coupling in bacteria and dependency between translation and mRNA stability has been established by seminal studies that used *lacZ* as a model gene^9,28,33,34^. Its regulatory mechanisms are well characterized, allowing us to manipulate parameters for *lacZ* gene expression and test hypotheses toward a generalizable model. For example, we introduced the effect of transertion to *lacZ* to emulate what happens to genes encoding inner membrane proteins^22^, we varied the 5’ untranslated region (5’-UTR) sequence of *lacZ* to test variable translation efficiencies across the genome^3,35^, and we perturbed the subcellular localization of RNase E to capture differences across bacterial species^36,37^. From this approach, we identified spatial and genetic design principles that bacteria have evolved to differentially regulate transcriptional and translational coupling to mRNA degradation across various genes and species.

## Results

### *lacZ* mRNA is degraded post-transcriptionally, uncoupled from transcription

While the possibility of nascent mRNA degradation during transcription has been discussed^12–15,27,38,39^, the actual rate of co-transcriptional mRNA degradation has never been reported. We used *lacZ* gene under the *lac* promoter in *E. coli* as a model to measure the rates of co-transcriptional and post-transcriptional mRNA degradation (*k*_d1_ and *k*_d2_, respectively; **Fig. 1A**). Earlier studies discussed co-transcriptional degradation of *lacZ* based on the observation that *lacZ* decays before the synthesis of *lacY* and *lacA* in the original lac operon. However, this result can be explained by co-transcriptional mRNA processing at the intergenic region between *lacZ* and *lacY*^40–42^, instead of real co-transcriptional degradation of *lacZ*. Therefore, we deleted *lacY* and *lacA* genes from the original *lac* operon in the chromosome of wild-type, MG1655 to yield a monocistronic *lacZ* (strain SK98). The intrinsic terminator sequence after *lacA* follows the coding sequence of *lacZ* to ensure the dissociation of RNA polymerase (RNAP) from DNA after finishing the transcription of *lacZ* (**Fig. 1B**). To follow the degradation kinetics of *lacZ* mRNA after the stoppage of transcription initiation, transcription of *lacZ* was induced with membrane-permeable inducer isopropyl b-D-1-thiogalactopyranoside (IPTG) and re-repressed with glucose 75 seconds (s) after addition of IPTG (**Fig. 1B**). Importantly, glucose was added before the first RNAPs finished transcription, so that co-transcriptional mRNA degradation can be observed. During this time course, a population of cells was acquired every 20-30 s, from which 5’ and 3’ *lacZ* mRNA levels were quantified by quantitative real-time PCR (qRT-PCR) using probes denoted as Z5 and Z3, respectively. A key feature of this time course experiment is the temporal separation of *lacZ* mRNA status between nascent and released (**Fig. 1B**). Until the first RNAPs finish transcription of *lacZ* (T_3’_), all *lacZ* mRNAs are expected to be nascent (time window i). After the last RNAPs finish transcription of *lacZ* at t_3’_, all *lacZ* mRNAs are released (time window iii). In between, nascent and released *lacZ* mRNAs co-exist (time window ii). Hence, we can measure the rates of co-transcriptional and post-transcriptional mRNA degradation by fitting Z5 level changes with an exponential decay function in the time windows i and iii, respectively.

**Figure 1.**
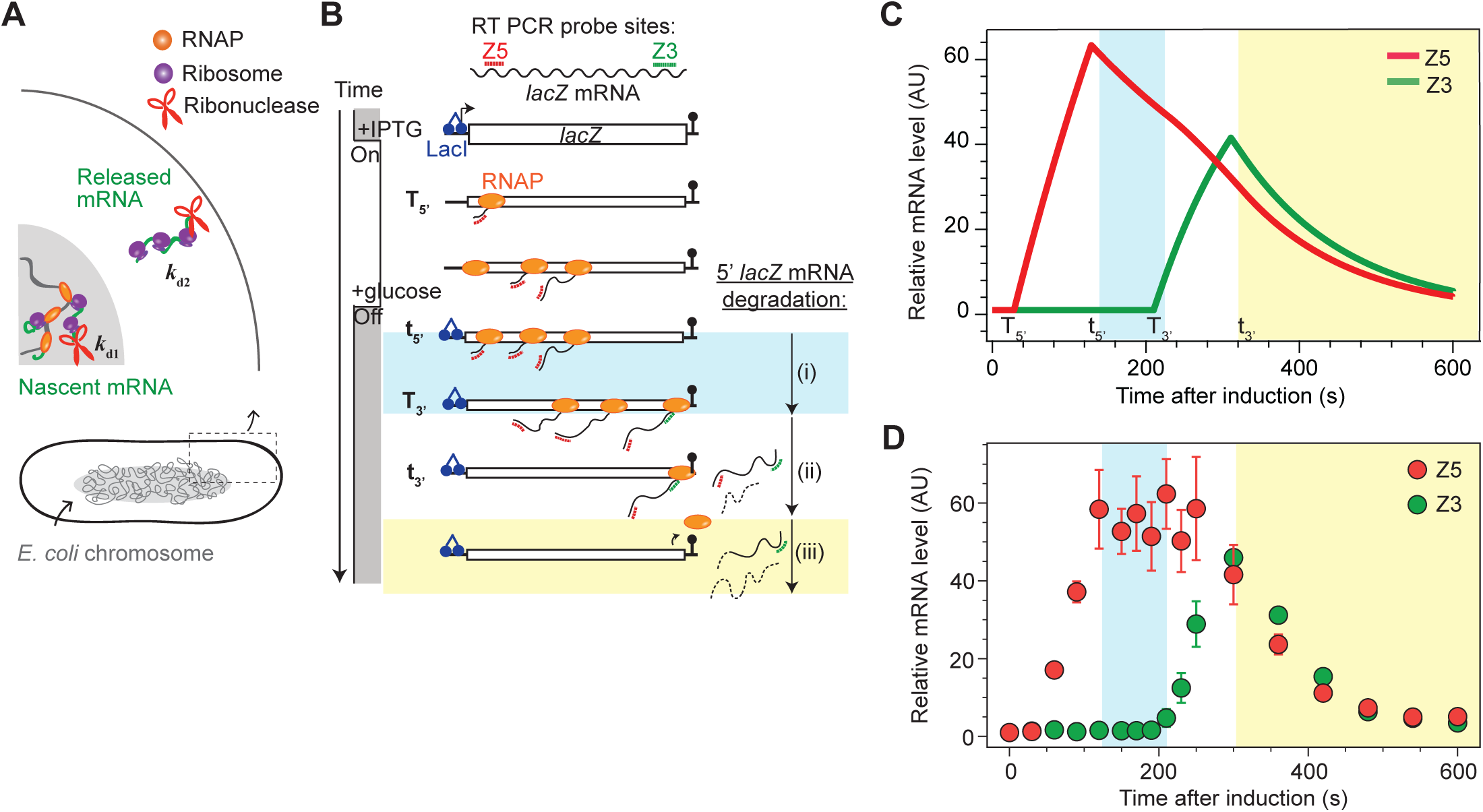
Two-phase mRNA degradation in bacteria. (**A**) Definition. mRNA degradation can occur during transcription (on nascent transcripts) and after transcription (on released transcripts) with different rates *k*_d1_ and *k*_d2_, respectively. (**B**) Schematics of the time-course assay, in which *lacZ* transcription is pulse-induced and its transcripts are quantified with qRT-PCR primers amplifying the 530-660 nucleotide (nt) region (Z5) and 2732-2890 nt region (Z3) of *lacZ* (gene length = 3072 nt). The first and last RNAPs pass the Z5 probe site at T_5’_ and t_5’_, respectively, and they pass the Z3 probe site at T_3’_ and t_3’_, respectively. Blue and yellow shaded boxes indicate the time when *k*_d1_ and *k*_d2_ can be measured by an exponential decay fit, respectively. (**C**) Anticipated result when mRNA degradation takes place with *k*_d1_ = 0.18 min^-1^ and *k*_d2_ = 0.42 min^-1^. (**D**) Time course data of 5’ and 3’ *lacZ* mRNA (Z5 and Z3) after induction with 0.2 mM IPTG at t = 0 and re-repression with 500 mM glucose at t = 75 s (strain SK98, grown in M9 glycerol at 30°C). Error bars represent the standard deviation from three biological replicates.

If the co-transcriptional degradation of *lacZ* mRNA takes place (*k*_d1_>0), Z5 level will decrease in the time window i, as demonstrated by a mathematical model (**Fig. 1C** and **S1A**). However, our data shows that Z5 level stays constant in the time window i, suggesting that *k*_d1_ is close to zero (**Fig. 1D**). From biological replicates, we obtained *k*_d1_ = 0.042 ± 0.0598 min^-1^ and *k*_d2_ = 0.43 ± 0.067 min^-1^ (**Fig. S1B**). Essentially, the mean lifetime of nascent *lacZ* mRNA (1/*k*_d1_) is 24 min, much longer than transcription elongation time (∼3.5 min), suggesting that *lacZ* mRNA is unlikely to experience degradation during transcription elongation.

### Membrane localization of RNase E accounts for uncoupling of transcription and degradation of *lacZ* mRNA

Among various ribonucleases in *E. coli*, the endoribonuclease RNase E has been considered the main enzyme to initiate mRNA degradation^21,36,43–45^, including *lacZ* mRNA^27,46,47^. To confirm that the observed *k*_d1_ and *k*_d2_ of *lacZ* mRNA are controlled by RNase E, we repeated the experiment in a strain carrying a temperature-sensitive RNase E allele (*rne*3071, strain SK519), in which RNase E can be inactivated by a 10-min shift to 43.5°C^48^. We performed IPTG induction of *lacZ* transcription at 43.5°C after 10 min of the temperature shift. Because transcription elongation is faster at this high temperature (T_3’_ = 100 s), glucose was added at 50 s after IPTG induction, so that we can still capture the time window i to measure *k*_d1_. When RNase E was inactivated, *k*_d1_ and *k*_d2_ were about 7 times smaller than those measured in wild-type RNase E at 43.5°C and about 2-3 times smaller than those measured at 30°C (**Fig. S2A**), confirming that RNase E controls *k*_d1_ and *k*_d2_ of *lacZ* mRNA. In *E. coli* cells, RNase E is associated with the inner membrane via the membrane targeting sequence (MTS)^20^. Therefore, the lack of co-transcriptional degradation of *lacZ* mRNA (very low *k*_d1_) could be accounted for by the membrane localization of RNase E, away from the nucleoid (or transcription site).

*E. coli* cells are viable even when the MTS sequence of RNase E is removed and RNase E is localized to the cytoplasm^20,49^ (RNase E ΔMTS, **Fig. 2A**). When RNase E is in the cytoplasm, instead of anchored to the membrane, it can interact with nascent and released mRNAs more frequently and likely affect *k*_d1_ and *k*_d2_ of *lacZ* mRNA. To check this possibility, we measured *k*_d1_ and *k*_d2_ of *lacZ* mRNA in the strain expressing RNase EΔMTS. We found that cytoplasmic RNase E increases both *k*_d1_ and *k*_d2_; especially, *k*_d1_ increases about 7 fold, to 0.31 ± 0.084 min^-1^, in comparison to the wild-type RNase E strain (**Fig. 2B**).

**Figure 2.**
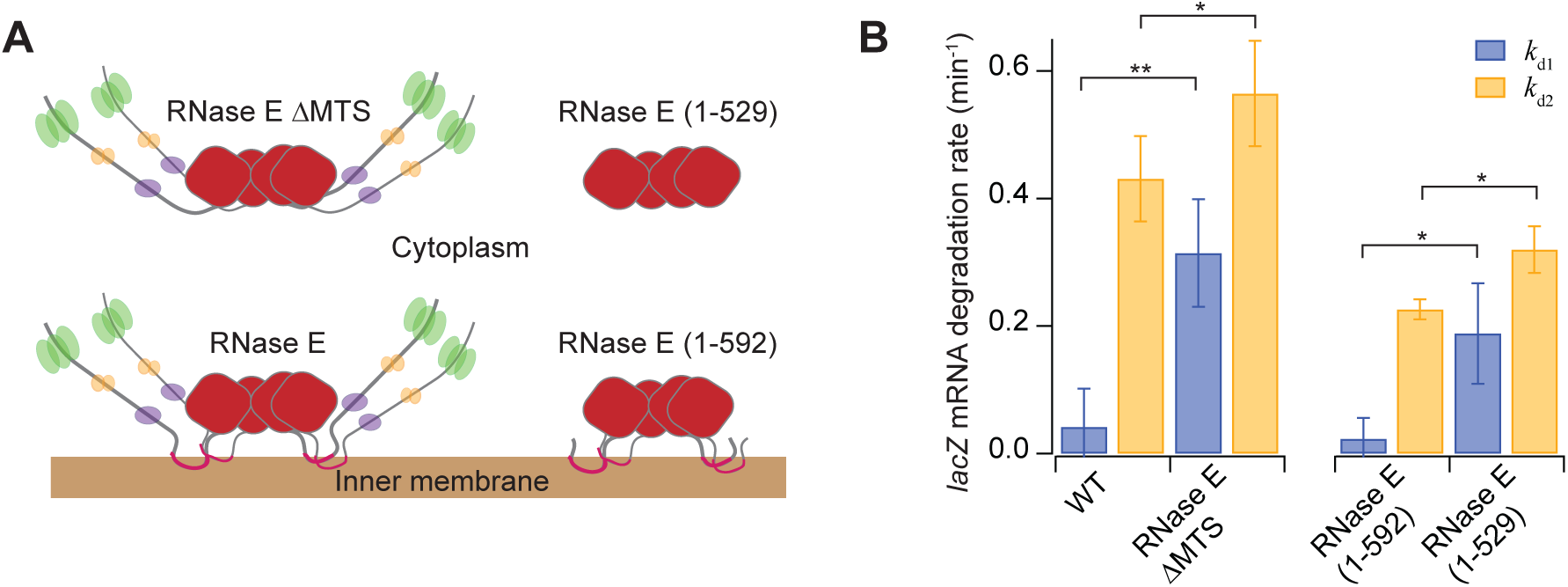
Effect of RNase E localization and its C-terminal domain on *k*_d1_ and *k*_d2_ of *lacZ* mRNA. (**A**) Schematic description of wild-type RNase E and mutants forming different RNA degradosome complexes. The wild-type RNase E interacts with PNPase (green), Enolase (yellow), and RhlB (purple) to form the RNA degradosome. Not drawn to scale. (**B**) *k*_d1_ and *k*_d2_ of *lacZ* mRNA in RNase E localization mutant strains (strain SK98, SK339, SK370, and SK369). Transcription of *lacZ* was induced with 0.2 mM IPTG at t = 0 and re-repressed with 500 mM glucose at t = 75 s. Error bars represent the standard deviation from three biological replicates. ** and * indicate p<0.01 and p<0.05, respectively (two-sample t test).

We tested another cytoplasmic RNase E mutant, RNase E (1-529) (**Fig. 2A**). This mutant lacks the MTS as well as the C-terminal domain, which provides binding sites for other RNA degradosome components^50^ (**Fig. S2B**). To interpret the effect of RNase E localization in the absence of the C-terminal domain, we compared *k*_d1_ and *k*_d2_ of *lacZ* mRNA from cells expressing the RNase E (1-529) mutant with those from cells expressing a RNase E (1-592) mutant, which also lacks the C-terminal domain but is localized to the membrane via MTS. We found that *k*_d1_ and *k*_d2_ of *lacZ* mRNA are higher in the cytoplasmic RNase E (1-529) than in the membrane-bound RNase E (1-592) (**Fig. 2B**), supporting that mRNA degradation is faster when RNase E is localized in the cytoplasm. We note that the absence of C-terminal domain in RNase E (1-592) results in lower *k*_d1_ and *k*_d2_ of *lacZ* mRNA in comparison to those in the wild-type RNase E, suggesting the importance of having the C-terminal domain for the catalytic activity of RNase E happening at the N-terminal domain^51,52^ (See **Fig. S2C** for additional data). Altogether, our results show that the membrane localization of RNase E slows down the degradation of *lacZ* mRNA, especially during transcription, giving rise to the uncoupling of transcription and mRNA degradation.

### Proximity of nascent mRNAs to the membrane alone does not affect their degradation rates

Since slow co-transcriptional mRNA degradation is likely due to the spatial separation between membrane-localized RNase E and nascent mRNAs, we considered a scenario where nascent mRNAs are positioned close to the membrane. When mRNAs coding for a transmembrane protein are transcribed, co-transcriptional translation may be accompanied by membrane insertion of the nascent protein, a process known as transertion^53–55^. A previous study showed that expression of *lacY* (encoding the lactose permease localized in the inner membrane) brings the *lacY* locus and nearby DNA region (∼90 kb) close to the membrane^23^. This suggests that even a gene encoding a cytoplasmic protein (such as *lacZ*) can be localized close to the membrane if it is adjacent to an actively transcribed *lacY* gene locus on the chromosome. Therefore, we inserted a constitutively expressed *lacY* gene downstream of *lacZ* (strain SK435; **Fig. 3A**) to test if the transertion of *lacY* can bring *lacZ* closer to the inner membrane and increase *k*_d1_. As a control, we made a strain where *lacY* is replaced with *aadA*, a gene encoding a cytoplasmic protein, spectinomycin adenylyltransferase, that does not undergo transertion (strain SK390).

**Figure 3.**
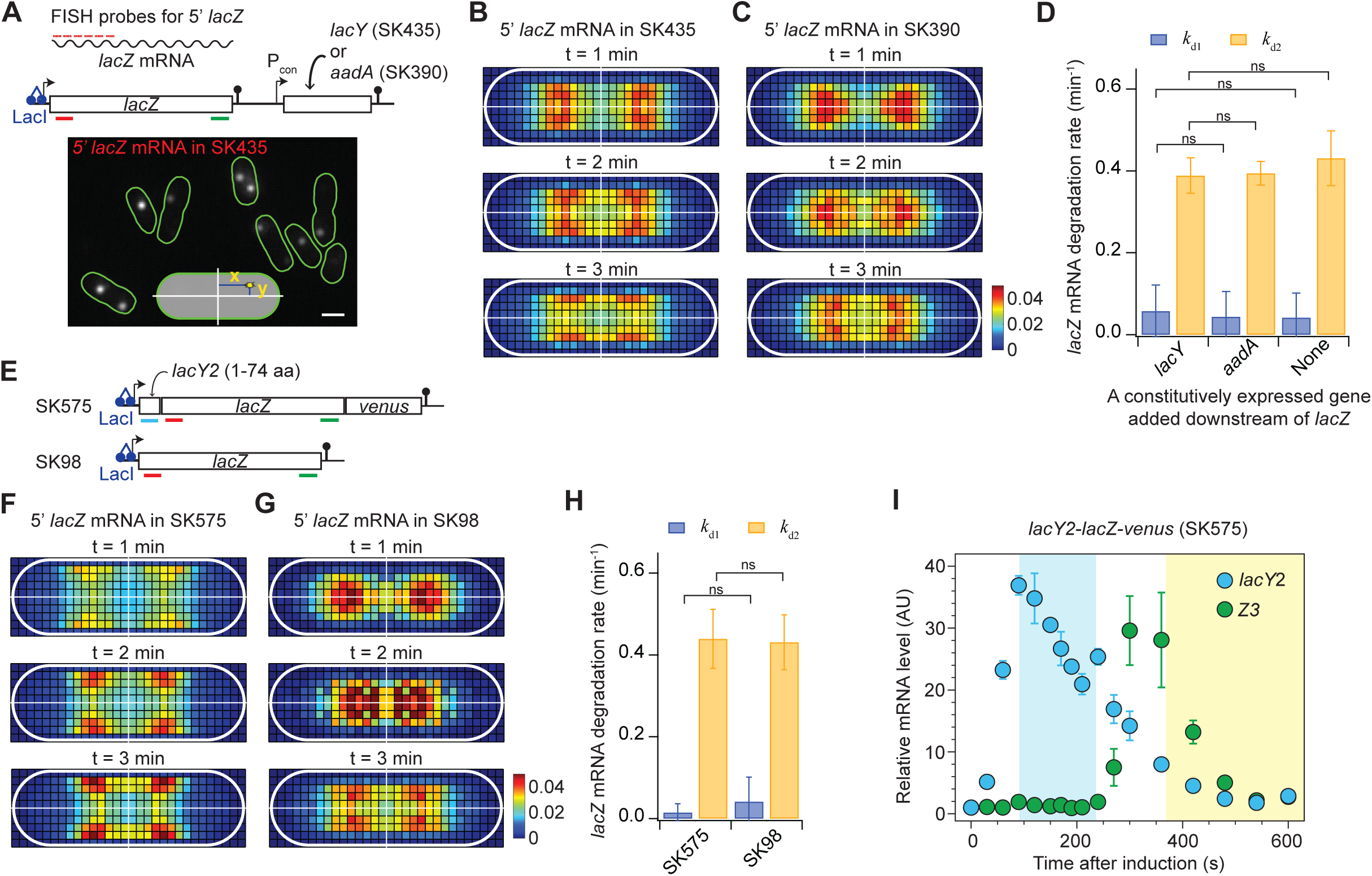
Effect of transertion on *k*_d1_ and *k*_d2_ of *lacZ* mRNA. (**A**) Localization of 5’ *lacZ* mRNA in the presence of active transcription of *lacY* or *aadA* from a constitutive promoter P_con_. An example FISH image of 5’ *lacZ* mRNA when transcription was induced with 0.2 mM IPTG and re-repressed with 500 mM glucose at 75 s after induction. The example image is from cells taken at t = 1 min. Scale bar = 1 µm. (**B**-**C**) 2D histogram of Z5_FISH_ localization at different time points along the time-course assay. Colors denote the probability of finding Z5_FISH_ in a certain bin location. To minimize noise, the normalized positions of foci along the cell long and short axes were calculated in the first quartile and extended to the other three quartiles using mirror symmetry. The bin size is 70-80 nm. The white ovals are cell outlines, and the white lines are the axes of symmetry. For each histogram, over 5,000 foci were analyzed. The same color scale was used for both histograms. (**D**) *k*_d1_ and *k*_d2_ of *lacZ* mRNA with different downstream genes: *lacY* (SK390), *aadA* (SK435), or none (SK98). (**E**) A pair of strains to study the direct transertion effect on *lacZ* mRNA degradation kinetics. (**F**-**G**) 2D histogram of Z5_FISH_ localization in *lacY2-lacZ-venus* (SK575, **F**) and *lacZ* (SK98, **G**). Transcription of *lacZ* was induced and re-repressed the same way as described in panel A. For each histogram, over 2,000 mRNA foci were analyzed. Except only 564 foci were used for SK98 t = 2 min. (**H**) *k*_d1_ and *k*_d2_ of *lacZ* mRNA in SK575 and SK98. (**I**) Relative mRNA level of the *lacY*2 sequence in SK575 measured by qRT-PCR during the time course assay described in panel A. *lacY*2 is probed by primers amplifying 80-222 nt region of *lacY*2 sequence. Blue and yellow shaded boxes indicate the time windows i and iii for *k*_d1_ and *k*_d2_ of *lacY*2, respectively. In all panels, error bars represent the standard deviation from three biological replicates. Also, ns indicates a statistically nonsignificant difference (two-sample t test).

To test the effect of transertion on the localization of nascent *lacZ* mRNA, we performed fluorescence in situ hybridization (FISH) using Cy3B-labeled probes binding to the first 1-kilobase region of *lacZ* mRNA (Z5_FISH_; **Fig. 3A**)^56^. Transcription of *lacZ* was induced with IPTG and re-repressed with glucose as in the qRT-PCR experiment (**Fig. 1B**). Cells were sampled every 1 min interval and fixed immediately. The Z5_FISH_ signal appeared as diffraction-limited foci (**Fig. 3A** and **S3A**). Their centroid coordinates along the short and long axes of the cell were normalized to cell width and length, respectively, and combined into a 2D histogram (**Fig. 3B**-**3C**). Notably, until T_3’_ (or the end of the time window i, t = 210 s), most of the Z5_FISH_ signals are expected to be from nascent mRNAs tethered to gene loci (**Fig. 1B**). Hence, the location of Z5_FISH_ at t = 1, 2, and 3 min after induction allows us to examine the subcellular localization of the nascent mRNAs (and their gene loci) exclusively. We observed that already at t = 1 min, Z5_FISH_ in SK435 (*lacZ* followed by constitutively transcribed *lacY*) were localized off the center long axis, while those in SK390 (*lacZ* followed by constitutively transcribed *aadA*) were close to the center long axis of the cell (**Fig. 3B**-**3C**). As time progresses to t = 2 and 3 min, Z5_FISH_ in both strains localized away from the center long axis, likely due to the *lacZ* gene locus moving to the periphery of the nucleoid upon induction—an effect previously observed in the *lacZ* locus in *E. coli*^57^. In all three time points, Z5_FISH_ in SK435 were localized closer to the membrane than those in SK390 (**Fig. 3B**-**3C**), suggesting that the transertion of *lacY*, which does not occur with *aadA*, results in the neighboring *lacZ* gene transcription taking place close to the inner membrane.

Next, we measured *k*_d1_ and *k*_d2_ of *lacZ* mRNA in SK435 and SK390 by qRT-PCR to check if the proximity to the membrane allows the nascent and released *lacZ* mRNAs to be degraded faster. We found that *k*_d1_ and *k*_d2_ of *lacZ* mRNAs were invariable between the two strains and almost identical to the original *lacZ*-only strain without *lacY* or *aadA* (**Fig. 3D**).

We wondered if nascent *lacZ* mRNAs in SK435 were not close enough to the membrane to facilitate their degradation. To bring nascent *lacZ* mRNAs even closer to the membrane, we constructed a translational fusion of *lacZ* with the first two transmembrane segments of *lacY* (*lacY*2), so that *lacZ* is directly linked to the transertion element (**Fig. 3E**). We also fused the *venus* gene at the 3’ end of *lacZ* sequence to verify the membrane localization of LacZ proteins by fluorescence imaging (strain SK575; **Fig. S3B**). FISH imaging of 5’ *lacZ* mRNA expressed from the *lacY*2-*lacZ-venus* fusion at t = 1, 2, 3 min after induction showed that as soon as two transmembrane segments (*lacY*2 of mRNA length 222 nt) are transcribed, the *lacZ* sequence is strongly enriched near the membrane in comparison to the original *lacZ* strain without the *lacY2* fusion (**Fig. 3F**-**3G** and more information in **Fig. S3C**). This result suggests that transertion takes place immediately after induction and brings nascent transcripts to the membrane. Also, the direct translational fusion of *lacY*2 element to *lacZ* placed the nascent *lacZ* mRNAs closer to the membrane in comparison to the previous *lacZ*-*lacY* context where the transertion effect came from the neighboring *lacY* gene locus (strain SK435; **Fig. 3B**).

While nascent *lacZ* mRNAs were closer to the membrane, their *k*_d1_ was not larger than that of the original *lacZ*-only strain (strain SK98; **Fig. 3H**). This result further supports the notion that proximity to the inner membrane (where RNase E is localized) is not sufficient to increase the rate of co-transcriptional *lacZ* mRNA degradation.

In the *lacY*2-*lacZ-venus* fusion strain, we can also measure the degradation kinetics of the *lacY*2 region of the transcripts using a set of qRT-PCR primers amplifying that region. Remarkably, we found that *lacY*2 exhibits fast co-transcriptional mRNA degradation with *k*_d1_ = 0.34 ± 0.041 min^-1^ (**Fig. 3I** and **S3D**). This likely represents the characteristics of the original *lacY* transcript, as we measured a similar rate of co-transcriptional mRNA degradation from the full-length *lacY* gene (**Fig. S3E**). This finding indicates that coupling between transcription and mRNA degradation is possible for transcripts encoding inner-membrane proteins. Considering that the proximity of nascent mRNAs to the membrane alone does not affect the co-transcriptional degradation rate of *lacZ* mRNA, the fast co-transcriptional degradation observed with *lacY* mRNA is likely due to additional factors other than its proximity to the membrane (see **Discussion**).

### Another cytoplasmic protein-coding gene, *araB*, exhibits high *k*_d1_ due to its RBS sequence

To determine if the result we observed for *lacZ* is generalizable to other genes in *E. coli*, we examined *k*_d1_ and *k*_d2_ of another gene encoding a cytoplasmic protein, *araB*, which is under the control of the arabinose-inducible promoter (P_ara_) of the *araBAD* operon on the chromosome of *E. coli* strain MG1655 (**Fig. 4A**). We deleted *araA* and *araD* genes to make *araB* a monocistronic gene (P_ara_-*araB*; strain SK472). Transcription from the P_ara_ promoter was induced with arabinose and re-repressed by glucose 50 s afterward. In contrast to very slow co-transcriptional degradation observed in *lacZ* mRNA, 5’ *araB* mRNAs were degraded before 3’ *araB* mRNAs were transcribed (time window i indicated as a blue box in **Fig. 4B**), resulting in *k*_d1_ = 0.55 ± 0.134 min^-1^ (**Fig. 4C**). This is quite striking because in *E. coli*, co-transcriptional degradation does not seem to occur for genes encoding cytoplasmic proteins due to the membrane localization of RNase E (**Fig. 2B**).

**Figure 4.**
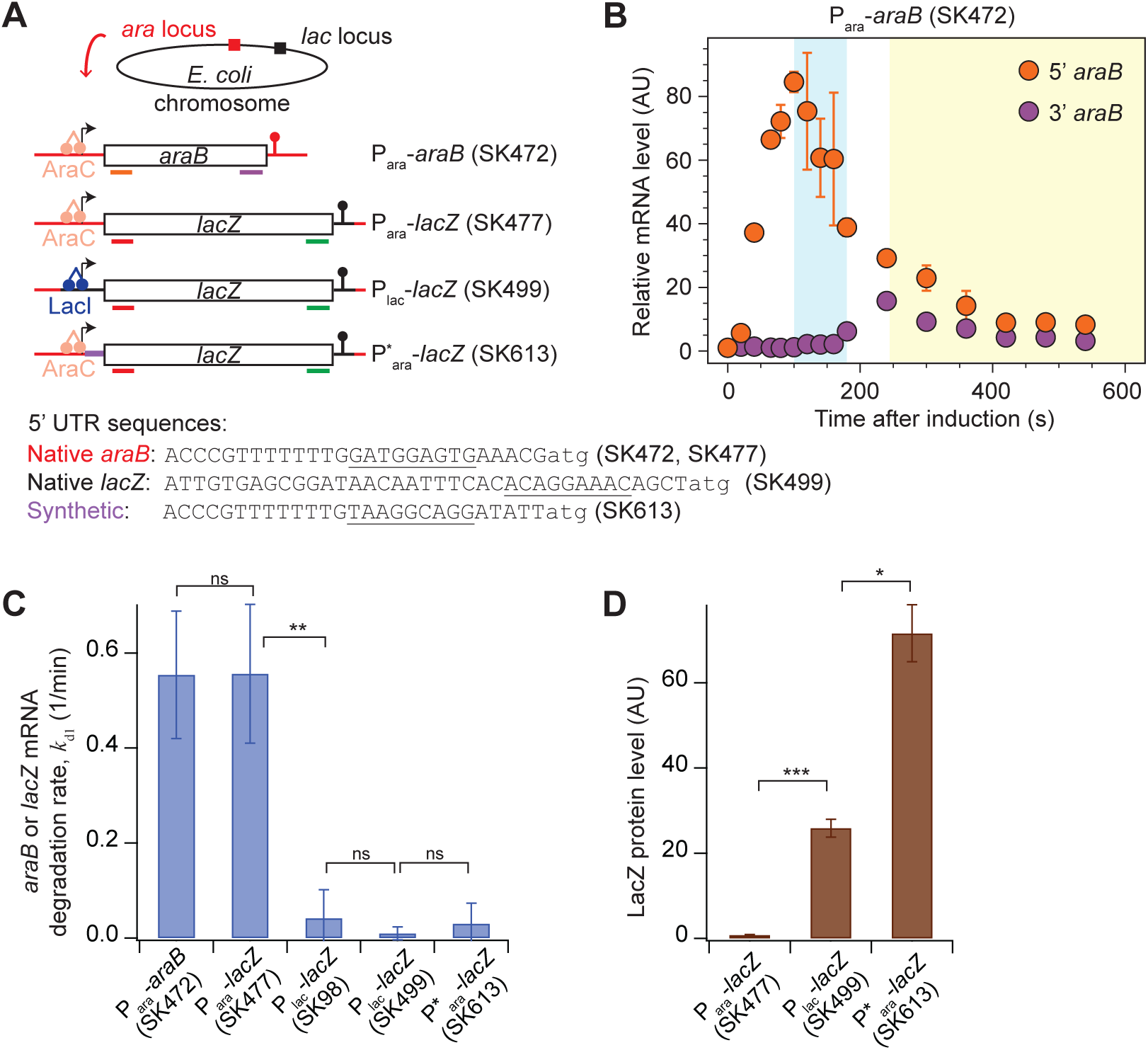
Degradation kinetics of *araB* mRNA and the effect of RBS sequences on *k*_d1_. (**A**) Design of strains used in this figure. 5’-UTR sequences from the first base (+1) of the transcript to the start codon (atg) are shown. SD elements estimated by an RBS calculator^63^ are underlined. We note that *lacZ* in the original *lac* locus was deleted when *lacZ* was placed in the *ara* locus. (**B**) *araB* mRNA level change from induction with 0.2% arabinose at t = 0 and re-repression with 500 mM glucose at t = 50 s. 5’ *araB* and 3’ *araB* were probed by qRT-PCR primers amplifying 33-210 nt and 1536-1616 nt regions of *araB*. Blue and yellow boxes denote the time windows where *k*_d1_ and *k*_d2_ are measured. (**C**) *k*_d1_ measured in strains shown in panel A. P_ara_ and P*_ara_ were induced with 0.2% arabinose, and P_lac_ was induced with 0.2 mM IPTG. 500 mM glucose was added at t = 50 s (for *araB*) or 75 s (for *lacZ*) to turn off the promoter. (**D**) LacZ protein expression measured by Miller assay. LacZ expression was induced and re-repressed the same way as in the qRT-PCR experiment (panel C), and the total LacZ protein produced from the pulsed induction were calculated from each strain. In all panels, error bars indicate the standard deviations from three or more biological replicates (except D, from two replicates). ***, **, and * indicate p<0.001, 0.01, and 0.05, respectively, and ns indicates a statistically nonsignificant difference (two-sample t test).

We found that this high *k*_d1_ is not due to any aspects of the *araB* sequence, because replacing *araB*’s coding region with that of *lacZ* (P_ara_-*lacZ*; SK477) resulted in similarly high *k*_d1_ of 0.56 ± 0.146 min^-1^ of *lacZ* mRNA, in contrast to the low *k*_d1_ observed at the native *lac* locus (SK98; **Fig. 4C**). The high *k*_d1_ of *lacZ* mRNA produced from P_ara_ was not due to the chromosomal position either, because bringing the *lacI-lacZ* region from the native *lac* locus to the *ara* locus (P_lac_-*lacZ*; SK499) did not change the original (low) *k*_d1_ of *lacZ* mRNA (SK98; **Fig. 4C**). The high *k*_d1_ likely originates from what P_ara_-*araB* (SK472) and P_ara_-*lacZ* (SK477) have in common: the sequence in 5’-UTR (**Fig. 4A**), which is different from 5’-UTR of native *lacZ* (SK98 and SK499). Therefore, the high *k*_d1_ of P_ara_ may originate from a certain feature of the 5’-UTR sequence.

5’-UTR of an mRNA contains ribosome binding site (RBS), including Shine-Dalgarno (SD) sequence, which governs translation initiation and protein expression level^58–60^. Henceforth, we refer to the SD sequence and its surrounding sequence as the RBS. RBS sequences are known to affect the energetics of ribosome binding and translation initiation, such that one can quantitatively predict the RBS strength, or protein expression outcome from the sequence^61–63^. However, weakening RBS strength by changing its sequence has also been known to destabilize the mRNA^27–31^, thus reducing the overall protein expression by reducing both translation initiation rate and mRNA lifetime.

Conforming to this expectation, *lacZ* transcripts with the native RBS of *araB* (P_ara_-*lacZ* in SK477) produced 30-fold lower LacZ protein expression than P_lac_-*lacZ* at the same chromosome location (**Fig. 4D**). This result came from measuring LacZ protein expression by Miller assay. To corroborate this finding, we replaced the RBS sequence in P_ara_-*lacZ* (SK477) with a strong RBS sequence designed using an RBS calculator^63^ (SK613 in **Fig. 4A**). The synthetic RBS sequence yielded increased LacZ protein expression, higher than that from native *lacZ* RBS (SK499; **Fig. 4D**). Also, *lacZ* mRNA with this strong synthetic RBS sequence exhibited low *k*_d1_ as observed in the native *lacZ* RBS (SK98 or SK499; **Fig. 4C**). These results support that the hypothesis that the weak RBS sequence in P_ara_-*araB* is responsible for the high *k*_d1_ of *araB* mRNA.

### RBS strength affects *lacZ* mRNA localization due to premature transcription termination

We have shown that P_lac_ and P_ara_, two inducible promoters widely used in gene expression studies^64–66^, have vastly different RBS strengths. Indeed, RBS sequences and their expected strengths vary widely among genes in the *E. coli* genome^3,35^. While mRNAs with a weaker RBS are expected to have shorter lifetime^16,17^, how RBS sequences affect co-transcriptional and post-transcriptional mRNA degradation rates has not been studied. To address this question, we compared the original *lacZ* strain (with the native RBS; SK98) with a weak RBS mutant, which was created by changing five bases in the original SD sequence (**Fig. 5A** and **S4A** for LacZ protein expression). We found that mutating the RBS sequence increases *k*_d1_ by 15 fold to 0.65 ± 0.171 min^-1^ without affecting *k*_d2_ (**Fig. 5B**). We confirmed that the high *k*_d1_ is largely controlled by RNase E because the temperature-sensitive RNase E allele (*rne*3071) showed much lower *k*_d1_ for this weak RBS at the non-permissive temperature in comparison to the wild-type RNase E at the same temperature (**Fig. S4B**-**S4C**). This brings us to the next question: How does membrane-bound RNase E carry out co-transcriptional degradation of mRNAs with a weak RBS?

**Figure 5.**
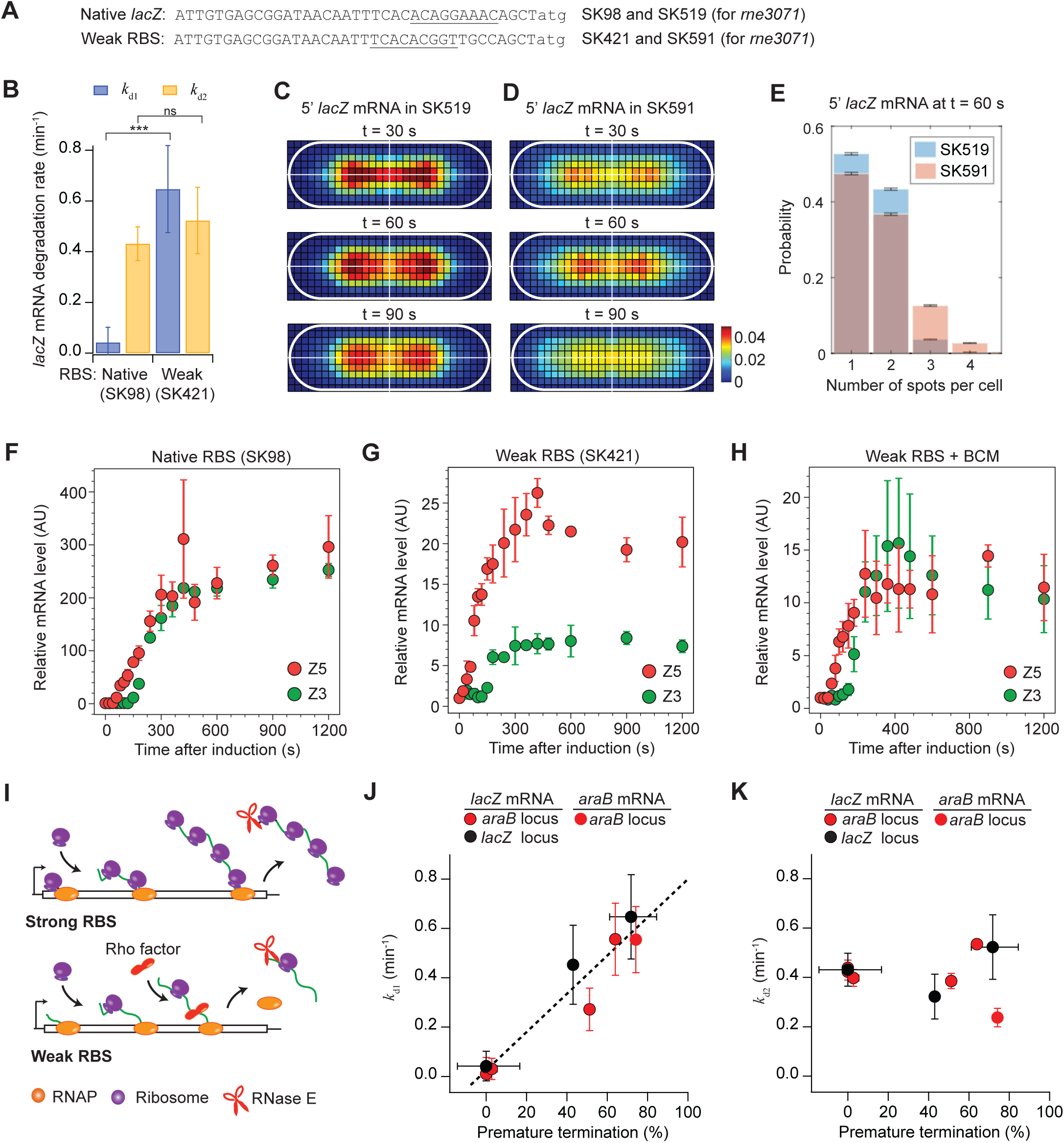
Origin of fast *k*_d1_ observed in *lacZ* mRNA with a weak RBS. (**A**) 5’ UTR sequences of native *lacZ* and a weak RBS mutant. mRNA sequences from the first base of the transcript to the start codon (atg) are shown. SD sequences estimated by an RBS calculator^63^ are underlined. (**B**) *k*_d1_ and *k*_d2_ of *lacZ* mRNA measured by induction with 0.2 mM IPTG at t = 0 and re-repression with 500 mM glucose at t = 75 s. Error bars represent the standard deviation from three biological replicates. *** denotes p<0.001, and ns indicates a statistically nonsignificant difference (two-sample t test). (**C-D**) 2D histogram of Z5_FISH_ localization depending on the RBS sequence. After shifting the temperature to 43.5°C for 10 min, *lacZ* expression was induced with 0.2 mM IPTG at t = 0 and re-repressed with 500 mM glucose at t = 50 s. In each case, over 25,000 mRNA foci were analyzed. (**E**) Number of fluorescent Z5_FISH_ spots detected per cell at t = 60 s during the time-course experiment described in panel C-D. Error bars represent the standard error from bootstrapping. (**F-H**) Z5 and Z3 levels after *lacZ* transcription was induced with 0.2 mM IPTG at t = 0. In (H), 100 µg/mL BCM was added 5 min before IPTG addition. Error bars represent the standard deviation from two biological replicates. (**I**) Effect of RBS strength on the fate of mRNA. (**J-K**) Relationship between *k*_d1_ or *k*_d2_ and the probability of premature transcription termination in various RBS-*lacZ* mRNAs and P_ara_-*araB* mRNA. See **Table S4** for the list of strains used. The line fit is based on the equation (1). Error bars for *k*_d1_ or *k*_d2_ represent the standard deviation from three replicates and those for *PT* were calculated from the steady-state ratio of Z5 and Z3 in two replicates.

To test the possibility that nascent mRNAs are localized differently depending on the RBS strength, we visualized 5’ *lacZ* mRNAs by FISH (Z5_FISH_). We reasoned that nascent mRNAs with a weak RBS sequence would be difficult to detect by FISH because they are quickly degraded (**Fig. 5B**). Therefore, we performed FISH in strains harboring the *rne*3071 allele to inactivate RNase E (strain SK519 and SK591 for the native and weak RBS sequences, respectively). At the non-permissive temperature, *lacZ* expression was induced with IPTG and re-repressed with glucose at 50 s after the induction. qRT-PCR analysis of RNA samples from this time-course experiment showed that Z3, probing the 3’ end of the mRNA, appears above the basal level at t = 100 s after induction (**Fig. S4C**), indicating that before t = 100 s, all 5’ *lacZ* mRNAs would be nascent and visualized as diffraction-limited foci originating from the gene loci that they are tethered to. Surprisingly, the 2D histogram of the relative positions of Z5_FISH_ in this time window showed different mRNA localization patterns between native RBS and weak RBS strains (**Fig. 5C**-**5D**). While Z5_FISH_ signals from the native RBS were localized at a specific location with a high probability (red bins in the histogram) as seen earlier in WT RNase E (**Fig. 3G**), Z5_FISH_ signals from the weak RBS were localized at random places throughout the cytoplasm, such that the dense region (red color) did not show up in the histogram (**Fig. 5D**). Additionally, before t = 100 s, the weak RBS strain contained a higher number of Z5_FISH_ spots per cell that have weaker fluorescence intensity in comparison to those in the native RBS strain (**Fig. S4D**-**S4E**). For example, at t = 60 s, there are up to two *lacZ* gene loci per cell (**Fig. S4F**), but the weak RBS strain had 16% of cells with 3 or more Z5_FISH_ spots per cell, in contrast to 4% observed in the native RBS strain (**Fig. 5E**). These results are consistent with a scenario, in which 5’ *lacZ* mRNAs with the weak RBS become physically separated from gene loci even when all of them are expected to be tethered to the gene loci and form only one or two diffraction-limited fluorescence spots per cell (**Fig. S4F**).

The spatial dispersion of mRNAs with the weak RBS in the time window i is reminiscent of premature transcription termination, previously shown to follow transcription-translation uncoupling due to nonsense mutation, antibiotic treatment, and amino acid starvation^33,67–69^. To check the possibility of premature RNAP termination in our weak RBS construct, we examined Z5 and Z3 levels at steady state after induction. In the native RBS strain (SK98), Z5 and Z3 levels were equal at the steady state (**Fig. 5F**). Considering that Z5 and Z3 have equal lifetimes (**Fig. S1B**), the equal steady state level means that 100% of RNAPs that passed the Z5 probe region reached the Z3 probe region at the end of the gene, i.e., 0% premature transcription termination. However, in the weak RBS *lacZ* strain (SK421), we observed that the steady state level of Z3 is about half of that of Z5, indicating significant premature transcription termination (**Fig. 5G**). We confirmed that this premature transcription termination is controlled by the rho factor because treatment with bicyclomycin (BCM) rescued the Z5 and Z3 difference, bringing the steady-state Z5 and Z3 levels to equal in the weak RBS strain (**Fig. 5H**).

These results are consistent with the hypothesis that transcription-translation coupling requires a strong RBS, which allows the loading of the first ribosome to the RBS as soon as the RBS sequence is transcribed by an RNAP. In the case of a weak RBS, in which the first ribosome loading event is delayed, an RNAP might not be coupled with a ribosome and experience premature termination by the rho factor^68,70^ (**Fig. 5I**). Then, the prematurely released (short) transcripts may diffuse to the membrane and get degraded by RNase E on the inner membrane.

### Translation affects mRNA degradation via premature transcription termination, not via ribosome protection

A notable lesson from the weak RBS strain is that there are prematurely released transcripts in the time window i, in which we measured *k*_d1_ assuming all transcripts are nascent. Hence, the high *k*_d1_ observed in weak RBS strains (including the ones observed with *araB*’s RBS in **Fig. 4C**) likely includes the degradation of prematurely released mRNAs and is not a true rate of co-transcriptional mRNA degradation. To address this problem, we modeled *k*_d1_ as a weighted average of real co-transcriptional degradation rate of nascent mRNAs (*k*_d1*_) and post-release degradation rate of prematurely terminated mRNAs (*k*_dPT_):

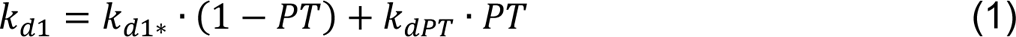

where *PT* is the probability of premature termination during transcription.

If premature transcription termination leads to high *k*_d1_ in a weak RBS strain, we expect to see a good correlation between *k*_d1_ and *PT*. To test this prediction, we used nine strains harboring *lacZ* with varying RBS sequences at the *ara* and *lac* loci (see **Table S4**). We measured *k*_d1_ and *k*_d2_ of *lacZ* mRNAs from re-repression (with glucose addition; **Table S5**) and calculated *PT* from steady-state levels of Z5 and Z3 after induction (without glucose addition). Because the ratio between stead-state levels of Z5 and Z3 (e.g. **Fig. 5F**-**5H**) is related to *PT* as well as *k*_dPT_ (**Fig. S5A**), we performed iterative fitting of *k*_d1_ and estimated *PT* using equation (1) and obtained best *PT* value for each strain and *k*_dPT_ common among nine strains (**Fig. 5J**; see **method details**). The optimal fitting of equation (1) gave *k*_d1*_ of 0.025 ± 0.0372 min^-1^ and *k*_dPT_ of 0.80 ± 0.0587 min^-1^ (*R*^2^ = 0.93). We note that *k*_d1*_ value is very similar to *k*_d1_ of strong RBS cases (where premature termination is 0%; e.g. SK98), and *k*_dPT_ value is larger than most of *k*_d2_ we have observed for transcripts released after transcription is completed. Possibly, prematurely released transcripts are degraded faster because they diffuse faster than longer, full-length mRNAs and/or because they lack certain features at the 3’ end that full-length mRNAs have, such as a stem-loop structure, making them more easily degraded, by 3’-to-5’ exonuclease, PNPase.

One of the models explaining the RBS effect on mRNA lifetime is based on the notion that ribosomes protect mRNA from the attack of RNase E^16,17^. According to this model, transcripts with a weak RBS sequence, or those showing high probability of premature transcription termination (*PT*), would undergo fast degradation because there are fewer ribosomes on the mRNAs. To test this model across different RBS sequences, we examined *k*_d2_, the decay rate of Z5 after t_3’_ (last RNAP passes the end of *lacZ* gene) in time window iii. *k*_d2_ is largely determined by the degradation rate of full-length transcripts that have the 3’ sequence and not affected by prematurely released transcripts, which are degraded rather quickly (*k*_dPT_) and minimally contribute to the Z5 signal in this time window. Nine strains of varying RBS sequences showed that *k*_d2_ is independent of *PT* (**Fig. 5K**) with very little correlation (P = -0.078). This result is in contrast to what would be expected from the ribosome protection model, which would expect higher *k*_d2_ in transcripts with weaker RBS, or higher probability of premature transcription termination, because the transcripts carry fewer ribosomes on average. Therefore, our results suggest that translation affects mRNA lifetime mainly by affecting the percentage of prematurely released transcripts (**Fig. 5J**).

### Premature transcription termination and subcellular localization of RNase E (or its homolog) affect the degradation of *lacZ* mRNA in other bacterial species

Our results so far imply that in *E. coli*, the fate of mRNA is determined by the RBS sequence because of its effect on transcription-translation coupling. Next, we examined if this conclusion can be generalized to other bacterial species. For example, in *B. subtilis*, RNAP was shown to translocate faster than the ribosome during expression of *lacZ,* preventing the ribosome from coupling to RNAPs^34,71^.We tested if the transcription-translation uncoupling in *B. subtilis* results in premature transcription termination and potentially a high *k*_d1_. First, we repeated the experiment done by previous papers measuring the transcription and translation times of *lacZ* by qRT-PCR and Miller assay in *B. subtilis* (strain GLB503; **Fig. 6A**), respectively. The transcription time was acquired from the initial increase of Z3 signal from the baseline after induction with IPTG, indicating the moment first RNAPs reach the end of the gene. The translation time was acquired from the initial increase of LacZ protein levels from the baseline after induction, indicating the moment first ribosomes reach the end of *lacZ* mRNA. In a slow growth condition^34^, we observed that the translation time was 2.6 ± 0.054 min, much longer than the transcription time of 1.3 ± 0.56 min (**Fig. S6A**-**S6C**). The steady-state levels of Z5 and Z3 were similar (**Fig. 6B**), implying that premature transcription termination does not take place even if transcription and translation are uncoupled in *B. subtilis*.

**Figure 6.**
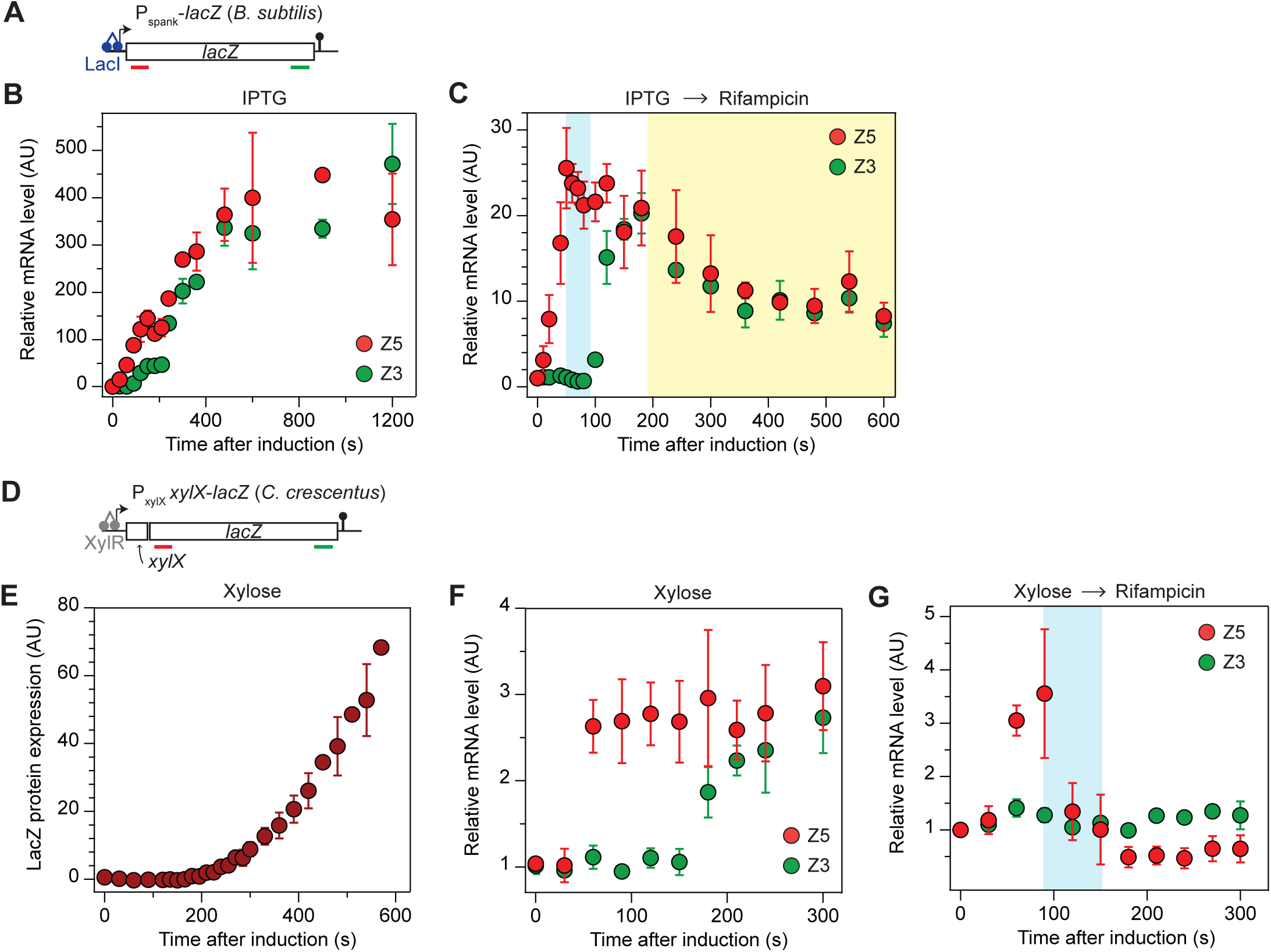
Degradation kinetics of *lacZ* mRNA in *B. subtilis* and *C. crescentus*. (**A**) IPTG-inducible *lacZ* in the chromosome of *B. subtilis*. For qRT-PCR, we used the same Z5 and Z3 primers used in *E. coli lacZ*. (**B-C**) Z5 and Z3 levels after induction with 5 mM IPTG at t = 0, probed by qRT-PCR. To measure *lacZ* mRNA degradation rates in (**C**), transcription was re-repressed with 200 µg/mL rifampicin at t = 30 s. The time windows used for *k*_d1_ and *k*_d2_ fitting are indicated as blue and yellow boxes. *B. subtilis* cells were grown in MOPS media supplemented with maltose at 30°C. Error bars represent the standard deviation from two (B) or three (C) biological replicates. (**D**) Xylose-inducible *lacZ* in *C. crescentus*. For qRT-PCR, we used the same Z5 and Z3 primers used in *E. coli lacZ*. (**E**) Translation kinetics of LacZ protein expression in *C. crescentus* after adding 0.3% xylose, probed by Miller assay using MUG (3-methylumbelliferyl-beta-D-galactopyranoside) as a sensitive LacZ substrate. Error bars represent the standard deviation from three biological replicates. (**F-G**) Z5 and Z3 levels after induction with 0.3% xylose at t = 0, probed by qRT-PCR. To measure *lacZ* mRNA degradation rates in (**G**), transcription was re-repressed with 200 µg/mL rifampicin at t = 50 s. The time window used for on *k*_d1_ fitting for Z5 is indicated as blue box. *C. crescentus* cells were grown in M2G at 28°C. Error bars represent the standard deviation from five (F) or three (G) biological replicates.

Next, we measured *k*_d1_ and *k*_d2_ of *lacZ* mRNAs by re-repressing transcription by adding rifampicin, a drug that stops transcription initiation^72–74^, at t = 30 s after induction (**Fig. 6C**). We obtained *k*_d1_ of 0.025 ± 0.0036 min^-1^ and *k*_d2_ of 0.14 ± 0.026 min^-1^ (**Fig. S6D**). Since premature transcription termination is not observed (**Fig. 6B**), *k*_d1_ can be attributed to co-transcription degradation, and the lifetime of nascent mRNA can be estimated as 40 min (1/*k*_d1_), much longer than the transcription time of 1.3 min. Hence, co-transcriptional degradation of *lacZ* mRNAs is likely very rare in *B. subtilis*, like in *E. coli*. The lack of co-transcriptional degradation can be explained by the membrane localization of the main endoribonuclease performing mRNA degradation, RNase Y and RNase E, in *B. subtilis* and *E. coli*, respectively^18,75,76^. We note that *k*_d2_ in *B. subtilis* is low relative to *E. coli* (see **Fig. 5K**), and *lacZ* mRNA levels do not return to the basal level within 10 min (**Fig. 6C**). This is quite surprising because the amount of LacZ proteins expressed from the 30-s induction was minimal according to the Miller assay using a sensitive fluorogenic LacZ substrate (**Fig. S6E**), suggesting that the remaining *lacZ* mRNAs do not support protein synthesis. Possibly, translation initiation is slow in this strain, and/or functional inactivation of mRNAs precedes the chemical degradation of mRNAs in *B. subtilis*.

In contrast to *E. coli* and *B. subtilis*, *C. crescentus* is known to have cytoplasmic RNase E^24–26^. Hence, we investigated the possibility of co-transcriptional degradation of *lacZ* mRNA in *C. crescentus* using a strain where *lacZ* is placed under the xylose-inducible promoter in the chromosome (strain LS2370^77^; **Fig. 6D**). First, we measured the transcription and translation times of *lacZ* by qRT-PCR and by Miller assay after induction with xylose. The translation time was 2.5 ± 0.21 min (**Fig. 6E** and **S6F**), similar to the transcription time of 2.3 ± 0.27 min (**Fig. 6F**), suggesting that transcription and translation are coupled. To measure *k*_d1_ and *k*_d2_ of *lacZ* mRNAs, transcription was re-repressed with rifampicin at t = 50 s after addition of xylose. qRT-PCR data show that Z5 decays very quickly after rifampicin addition (**Fig. 6G**). Strikingly, Z3 does not increase above the basal level, such that the time windows i and iii cannot be defined for fitting Z5 for *k*_d1_ and *k*_d2_. If we take T_3’_ of 2.3 min from the induction-only experiment (**Fig. 6F**) to estimate the time window i (blue box in **Fig. 6G**), we obtain *k*_d1_ of 1.3 ± 0.13 min^-1^. The absence of 3’ *lacZ* mRNA signals in the re-repression experiment (also minimal LacZ protein expression; **Fig. S6G**) indicates significant premature transcription termination, which contributes to high *k*_d1_.

Indeed, when we blocked the rho factor activity with BCM, the steady-state levels of Z5 and Z3 increased, suggesting that rho-dependent premature termination affected *lacZ* mRNA levels in non-treated cells (**Fig. S6H** vs. **6F**). Repeating *k*_d1_ measurement in BCM-treated cells allowed us to obtain the true rate of co-transcriptional degradation, *k*_d1*_ of 0.71 ± 0.093 min^-1^ (**Fig. S6I**). Through mathematical modeling, we estimated that prematurely terminated mRNAs are degraded at *k*_dPT_ of 3.4 ± 0.61 min^-1^ and the probability of premature transcription termination (*PT*) to be 69 ± 4.4% (see **method details**). The fast mRNA degradation likely originates from the cytoplasmic distribution of RNase E in *C. crescentus* cells. Interestingly, RNase E in *C. crescentus* has been shown to interact with the rho factor^78^. We speculate that the cooperation between the rho factor and RNase E results in the high *k*_d1_ and high probability of premature transcription termination we observed in *C. crescentus* (**Fig. 6G**).

The high probability of premature transcription termination (∼69%) agrees with the absence of 3’ *lacZ* mRNA signal when *lacZ* transcription was induced only for 50 s (**Fig. 6G**). However, it seems incompatible with transcription-translation coupling concluded based on the synchronized transcription and translation times (**Fig. 6E**-**6F**). We note that the transcription and translation times were determined by the first (fastest) RNAPs and ribosomes arriving at the 3’ end, and they can be the same even though only a small fraction of RNAPs are coupled to a ribosome. Hence, it is likely that a significant fraction of RNAPs is uncoupled with a ribosome during the transcription of *lacZ* in *C. crescentus* and experiences premature transcription termination.

## Discussion

Our findings have implications for gene regulation based on when and where mRNAs are degraded within a bacterial cell. In bacteria, such as *E. coli* and *B. subtilis*, where major ribonuclease and RNA degradosome are localized to the membrane, co-transcriptional mRNA degradation is likely negligible for most genes, and mRNA degradation takes place exclusively on the membrane once mRNAs are released from the gene loci (**Fig. 7A**-**7B**). The lack of co-transcriptional degradation would be advantageous when more proteins need to be made per transcripts.

**Figure 7.**
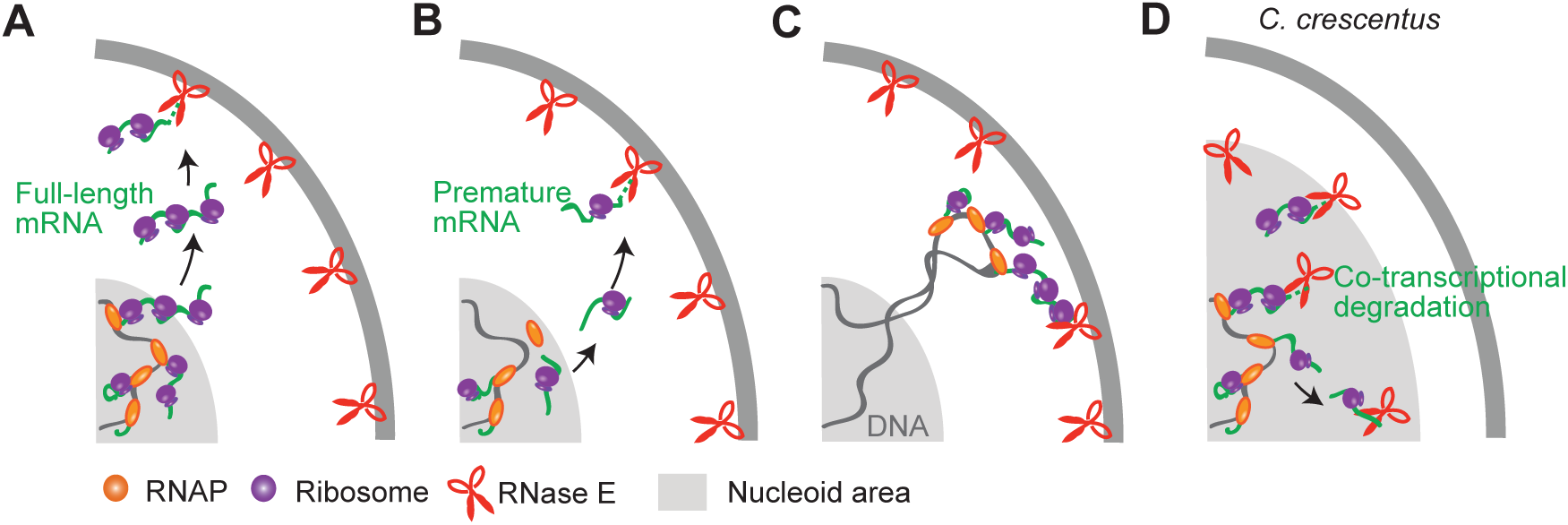
Generalizable model of mRNA degradation in bacteria. (**A**-**C**) Scenarios in *E. coli* (and possibly other bacterial species having the main ribonuclease on the membrane) for genes encoding cytoplasmic proteins with strong RBS (**A**) and weak RBS (**B**) and for genes encoding inner membrane proteins (**C**). (**D**) A scenario in *C. crescentus* and possibly other bacterial species having the main ribonuclease in the cytoplasm. The cartoon is drawn to reflect that nucleoid takes a large area of the cytoplasm in *C. crescentus*^97^.

Our data showing co-transcriptional degradation of *lacY* mRNA suggests an exception to this rule for genes encoding inner membrane proteins (**Fig. 7C**). We note that the high *k*_d1_ of 5’ *lacY* mRNA measured in *lacY*2-*lacZ*-venus (SK575; **Fig. 3I**) and in full *lacY* mRNA (SK564; **Fig. S3E**) likely reflects genuine co-transcriptional degradation, without premature transcription termination because (1) the native (strong) *lacZ* RBS was used and (2) full *lacY* transcript (SK564) showed 0% premature transcription termination (**Fig. S3F**). In terms of the mechanism, membrane localization of nascent mRNAs may not be the only reason that *lacY* mRNA is degraded co-transcriptionally. The *lacZ* sequence within *lacY2-lacZ-venus* mRNA had similar *k*_d1_ and *k*_d2_ as those of the *lacZ*-alone case (SK98) even though their localization (or the proximity to the membrane) were vastly different (**Fig. 3H**). Hence, we speculate that there are additional features in the *lacY2* sequence (i.e. the first two transmembrane segments) that promote its co-transcriptional mRNA degradation. For example, the signal recognition particle (SRP) and SecYEG, proteins involved in transertion of LacY^79,80^, might interact with RNase E to promote the degradation of *lacY*2 sequence. Although this idea remains to be tested, such a mechanism can also explain earlier results that translational fusion of a SRP signal peptide to a random gene decreases the transcript’s lifetime^81^ and that fast degradation of *ptsG* mRNA encoding transmembrane glucose transporter (IIBC^glc^) requires the transmembrane segment of its protein^82^. Also, this hypothesis predicts that the rate of co-transcriptional mRNA degradation of *lacY* would decrease when RNase E is localized in the cytoplasm. Indeed, previous genome-wide characterization of mRNA lifetimes in the RNase E ΔMTS strain showed that many genes encoding inner membrane proteins are preferentially stabilized in this cytoplasmic RNase E mutant^81,83^. These results suggest that membrane localization of RNase E is important for differential regulation of membrane protein expression in comparison to cytoplasmic proteins. Since membrane surface area is limited (more than the cytoplasmic volume)^84^ and since membrane channel proteins (such as LacY) have a higher activity cost when expressed^84,85^, tight regulation of membrane protein expression, by employing co-transcriptional degradation mechanism, is likely beneficial for cellular fitness.

Another important determinant of mRNA degradation is the timing that transcripts are released from the gene. This timing can vary depending on the gene length and RNAP speed. Also, in the case of polycistronic genes, mRNA processing in the intergenic region^40–42^ can release the promoter proximal gene first while promoter distal gene is being transcribed. Adding to this list, our work highlights the important role played by RBS sequences in permitting premature release of incomplete transcripts (**Fig. 5J**).

Based on our model (**Fig. 1A**), the mean mRNA lifetime in the steady state is affected by the degradation rates of nascent (*k*_d1*_), fully-transcribed (*k*_d2_), and prematurely-released (*k*_dPT_) transcripts because these three types of mRNA can have distinct degradation rates. If ribosomes indeed protect mRNA from degradation^16,17^, each of these rates may increase with lower RBS strength. However, our data suggests that these rates do not vary much among the RBS mutant strains we examined; instead, the portion of prematurely released transcripts varies significantly (**Fig. 5J**-**5K**), eventually yielding different protein output for different RBS sequences (**Fig. S5B**-**S5E**). Considering that premature transcription termination is a hallmark event in the absence of transcription-translation coupling^86,87^ (**Fig. 5I**), we identified RBS sequences as the key genetic feature that can modulate the probability of transcription-translation coupling and subsequently mean mRNA lifetimes across the genome.

The RBS sequences we tested cover a wide range of translation initiation strengths observed in the genome^88^ (**Fig. S5G**-**S5H**). If we compare the maximum translation initiation strength we observed non-zero percentage of premature transcription termination (strain SK420 and SK518 in **Fig. S5G**) with endogenous translation initiation strengths across the *E. coli* genome^88^, we estimate that at least 58% of all genes may experience some percentage of premature transcription termination due to compromised transcription-translation coupling (**Fig. 5I** and **S5H**). This estimation is consistent with the high percentage of 3’-end mRNAs detected at the 5’ UTRs and inside of genes in a recent *E. coli* transcriptome analysis^89^. Collectively, these results support that transcription-translation uncoupling arising from low translation initiation rate and the resulting premature transcription termination are likely common across the *E. coli* genome.

T7 transcription systems in *E. coli*, often used for bioengineering and synthetic biology field^90^, are known to experience transcription-translation uncoupling because T7 RNAP outpaces the host ribosome (8-fold speed difference^91^), yet T7 RNAP does not prematurely terminate^92^. Based on our model of mRNA degradation in *E. coli*, we predict that transcripts made by T7 RNAPs are degraded once transcription is completed, as opposed to experiencing co-transcriptional degradation as proposed previously^46^.

We found that gene expression in *B. subtilis* is analogous to the T7 system, such that premature transcription termination is negligible even though RNAP and ribosome are uncoupled (**Fig. 6B**). We note that a recent study showed that *B. subtilis* RNAPs can prematurely dissociate from DNA during transcription of *lacZ* in a rho-independent manner, especially when their speed is slow^71^. Hence, premature transcription termination may be possible under certain conditions and in certain genes that are under the control of riboswitches and attenuators^93^ and help down-regulate protein expression in *B. subtilis*.

In bacteria, such as *C. crescentus*, where major ribonuclease and RNA degradosome are located in the cytoplasm, mRNA degradation may start during transcription (**Fig. 7D**). We observed in *C. crescentus*, high rate of co-transcriptional degradation rate (*k*_d1*_) for *lacZ* mRNA and significant premature transcription termination (*PT* = 69%). This high premature transcription termination suggests that many RNAPs transcribing *lacZ* were not coupled to a ribosome and points out that the equality between transcription and translation times (**Fig. 6E**-**6F**) may not be a good indicator for the percentage of transcription-translation coupling. Together with the fact that the rho factor physically interacts with RNase E in *C. crescentus*^78^, our results imply that transcription, premature transcription termination, and mRNA degradation are highly coupled in the cytoplasm of *C. crescentus*.

In conclusion, our work overall identifies subcellular localization of RNase E (or its homologue) and premature transcription termination in the absence of transcription-translation coupling (arising from weak RBS sequences) as spatial and genetic design principles by which bacteria have evolved to differentially regulate transcriptional and translational coupling to mRNA degradation across genes and species. These principles will serve the basis for quantitative modeling of protein expression levels across the genome^24,81,94^ and for comprehending the subcellular localization patterns of mRNAs found for different genes and in different bacteria species^24,81,94^. In the future, it would be interesting to investigate whether our findings are relevant to the coordination of transcription, translation, and mRNA degradation in other contexts where there is a lack of membrane-bound microcompartments, such as archaea^95^, chloroplast^36^, and mitochondria^96^.

## Acknowledgements

We thank Drs. Christine Jacobs-Wagner, Gene-Wei Li, Jason Peters, Jared Schrader, Lucy Shapiro, and X. Sunney Xie for strains and materials, Dr. Nam Ki Lee for an RNA extraction protocol, and Dr. Ido Golding for allowing us to use his real-time PCR machine in the beginning of the project. We also thank Brooke Ramsey for helping with experiments, Laura Troyer for illustrations, Dr. Marie Bao for editorial assistance, and the members of the Kim lab for critical reading of the manuscript. This work was supported by NSF Center for Physics of Living Cells (1430124), NSF Science and Technology Center for Quantitative Cell Biology (2243257), NIH (R35GM143203), and Searle Scholars Program.

## Author contributions

Conceptualization, Se.K. and Sa.K.; Methodology and investigation, Se.K., Y.W., A.H., and Sa.K.; Writing and visualization, Se.K., Y.W., A.H., and Sa.K.; Supervision and funding acquisition, Sa.K.

