## Supplemental Information for "Re-defining how mRNA degradation is coordinated with transcription and translation in bacteria"

### Materials and Methods

#### Strains and plasmids

A complete list of strains and plasmids used in this study is provided in **Table S1**, and a brief description of their construction is provided under the table. All constructs made in this study were verified by sequencing the engineered DNA regions. All oligonucleotides used for strain construction were obtained from Integrated DNA Technologies (IDT), unless otherwise noted.

#### Strain growth conditions

For all gene expression measurements, cell cultures were prepared in two steps. First, colonies were inoculated in a liquid medium and grown for 8-9 hours. Then, it was diluted >10,000 fold in a fresh medium (same type as the initial culture) and grown overnight. Experiments were performed when the culture was in the exponential growth phase (optical density at 600 nm, $OD_{600}$, 0.2 for *E. coli* and *B. subtilis*; $OD_{660}$ 0.2 for *C. crescentus*).

All *E. coli* cells were grown in liquid cultures (20-30 mL) of M9 minimal medium (6 g/L dibasic sodium phosphate anhydrous, 3 g/L monobasic potassium phosphate, 0.5 g/L sodium chloride, 1 g/L ammonium chloride, 2 mM magnesium sulfate, and 0.1 mM calcium chloride) supplemented with 0.2% glycerol, 0.1% casamino acids, and 1 mg/L thiamine at 30°C using a water bath shaker. In this growth condition, all *E. coli* strains had a similar doubling time as the wild-type strain (SK98) at 95.0 ± 9.40 min, except RNase EΔMTS (SK339), which had a doubling time of 117.0 ± 27.0 min.

For strains containing the temperature-sensitive mutant of RNase E (*rne*3071), cell cultures were prepared the same way as the other *E. coli* strains and shaken at 30°C until $OD_{600}$ reached 0.2. Then, they were transferred to a second water bath shaker set at 43.5°C 10 minutes before induction. Gene expression measurements were done at 43.5°C.

*B. subtilis* cells were grown in MOPS minimal medium with maltose (1X MOPS mixture, 1.32 mM dibasic potassium phosphate, 0.2% glutamate, 0.1 g/L tryptophan, and 0.4% maltose) at 30°C. The doubling time was 92 ± 9.22 min.

*C. crescentus* cells were grown in M2G (0.87 g/L dibasic sodium phosphate anhydrous, 0.54 g/L monobasic potassium phosphate, 0.50 g/L ammonium chloride, 0.5 mM magnesium sulfate, 0.5 mM calcium chloride, 0.01 mM ferrous sulfate, and 0.2% glucose) at 28°C. The doubling time was 142 ± 3.33 min.

#### Induction and re-repression of *lacZ* expression

To induce the expression of *lacZ* under $P_{lac}$ in *E. coli*, IPTG was added to a final concentration of 0.2 mM. This event marks time zero (t = 0 s). A constant volume of cell culture was withdrawn thereafter at certain time intervals. To re-repress *lacZ* expression, we added 500 mM glucose at t = 75 s[1]. When bicyclomycin (BCM) was used, 100 µg/mL of BCM was added 5 min before IPTG.

The glucose addition time was chosen based on two factors: First, it should be more than 1 minute before $T_{3'}$, so that the time window i is sufficiently long for $k_{d1}$ fitting. Since we withdraw samples every 15-20 s, about 3-4 samples can be taken during time window i of 1 min for good $k_{d1}$ estimation. However, if glucose is added too early on, not much Z5 signal develops. Considering these two factors, we decided to add glucose at t = 75 s for *lacZ* expression in our experimental condition. This glucose addition time was varied for a different gene (*araB*) and growth conditions (e.g. for temperature-sensitive RNase E mutant).

To induce the expression of *lacZ* (and *araB*) under $P_{ara}$, 0.2% arabinose was used for induction instead of IPTG. To re-repress, 500 mM glucose was added at t = 75 s for *lacZ* and t = 60 s for *araB*. We note that glucose can stop transcription initiation at $P_{ara}$ because the CRP binding site exists in the promoter[2].

For *lacZ* expression in *B. subtilis*, 5 mM IPTG was added for induction (t = 0), and 200 ug/mL rifampicin was added to stop transcription initiation (t = 30 s). This rifampicin addition time was chosen based on the same considerations used for glucose addition time.

For *lacZ* expression in *C. crescentus*, 0.3% D-xylose was added for induction (t = 0), and 200 ug/mL rifampicin was added to stop transcription initiation (t = 50 s). When BCM was used, 100 µg/mL of BCM was added 5 min before xylose.

#### Total RNA extraction and quantitative real-time polymerase chain reaction (qRT-PCR)

At each time point, 0.4 mL of cell culture was withdrawn and immediately mixed with an equal volume of pre-cooled RNAlater Stabilization Solution (Life Technologies). Two or three samples were collected before induction (t = 0) to accurately measure the baseline (mRNA level before induction). After induction, 16-18 samples were collected at the indicated time points. Sampling intervals ranged between 10 s and 60 s depending on the number of time points needed to obtain $k_{d1}$ or $k_{d2}$ from fitting the data points.

Upon sampling, cell mixtures were incubated on ice for 20 min, and then 0.2 mL of pre-cooled media was added to dilute the high salt content of the RNAlater solution. After 2-min centrifugation at 10,000 x g, the supernatant was discarded. Cell pellets were resuspended in 100 µL of lysozyme solution (10 mg/mL of lysozyme in 10 mM Tris-HCl (pH 8) with 1 mM EDTA) and vortexed for 20 s. Also, 0.5 µL of 10% SDS was added to the samples and vortexed for 10 s.

Next, we used PureLink RNA Mini Kit (Life Technologies) to extract total RNA. We followed the manufacturer's protocol with some modifications. Specifically, we used on-column DNase digestion using PureLink DNase set (Life Technologies) after applying Wash Buffer I and incubated at room temperature for 15 min. After DNase reaction, Wash Buffer I was applied again before moving on to Wash Buffer II. For elution, 50 µL of RNase-free water was used. A typical concentration of RNA in the eluates was about 60-80 ng/µL (measured by a NanoDrop Spectrophotometer). 2 µL of eluates was used for each qRT-PCR reaction using KAPA SYBR FAST qPCR Master Mix (KAPA Biosystems) and CFX Connect Real-Time System (Bio-Rad). Primer sequences are provided in **Table S2**.

Relative mRNA levels (e.g., **Fig. 1D**) were calculated from the fold change in mRNA levels relative to the baseline (pre-induction samples), or $2^{-\Delta Ct}$. Here, Ct stands for cycle threshold when the fluorescent signal passes a threshold. ΔCt was calculated by subtracting each sample's Ct value with the average Ct of pre-induction samples.

The times that 5' and 3' probe signals appear above the baseline (T5' and T3') indicate when the first RNAPs pass 5' and 3' probe regions, respectively. These times were obtained by linear fitting the initial rise of the 5’ and 3’ signal and finding their intersection point with their baselines.

The rates of mRNA degradation were obtained by linear fitting of log-scale relative mRNA levels within a certain time window. For example, for data shown in **Figure 1D**, Z5 values between t = 150 s and 210 s were used to calculate co-transcriptional degradation rate ($k_{d1}$), and Z5 values between t = 300 s and 600 s were used to calculate post-transcriptional degradation rate ($k_{d2}$). Z3 values between t = 300 s and 600 s can be used to calculate post-transcriptional rate of the 3’ mRNA ($k_{d2}$ of Z3), but we focused our analysis on the degradation rate of Z5 (or the 5’ end of the mRNA). The degradation rates of Z3 are shown in **Figure S1**, **S5**, and **S6**. The time window for fitting was adjusted for certain strains and growth conditions depending on $t_{5'}$ (when the last RNAPs pass the 5’ probe region), $T_{3'}$, and $t_{3'}$ (when the last RNAPs pass the 3’ probe region), which can be identified from the time when Z5 reaches maximum, when Z3 starts to appear from the baseline, and when Z3 reaches maximum, respectively.

**Miller assay**

We followed a previously described procedure[1] with modifications. At each sampling time, 900 µL of cell culture was withdrawn and immediately mixed with pre-cooled stop solution. The stop solution was 100 µL of 5 mg/mL chloramphenicol for *E. coli* and *C. crescentus* and 100 µL of 10 mg/mL chloramphenicol and 10 mg/mL erythromycin for *B. subtilis*. Cells were washed by centrifugation at 7,000 x g for 1 min (*E. coli*), 18,200 x g for 4 min (*B. subtilis*), or 6,000 x g for 4 min (*C. crescentus*) at 4°C, and cell pellets were resuspended in 340 µL of 1x Z buffer (30 mM disodium phosphate, 10 mM monosodium phosphate, 5 mM potassium chloride, 0.5 mM magnesium sulfate) containing the same concentration of chloramphenicol (and erythromycin) as before.

To measure the relative number of cells in each sample, $OD_{600}$ was measured in a microplate reader (Synergy HTX multi-mode reader, BioTek) using 190 µL of (washed) cells. 50 µL of lysis buffer (1x Z buffer with 0.0073% sodium dodecyl sulfate and 1.28% 2-mercaptoethanol) and 20 µL of chloroform were added to the remaining 150 µL of cells and vortexed for 10 seconds for lysis. Samples were incubated for 10 minutes and centrifuged at 150 x g for 10 seconds to remove large cell debris.

For *E. coli* strains, 120 µL of cell lysate and 30 µL of ortho-nitrophenyl-β-galactoside (ONPG) solution (4 mg/mL in the lysis buffer) were mixed in a 96-well plate, and $OD_{420}$ and $OD_{550}$ were measured overnight in the plate reader. LacZ activity was calculated using the following formula:

$$\text{Activity} = 1000 \cdot \frac{(\text{OD}_{420} - OD_{420,background}) - 1.75 \cdot (\text{OD}_{550} - OD_{550,background})}{\text{Reaction time} \cdot (\text{OD}_{600} - OD_{600,background})} \cdot \text{dilution factor} \qquad \text{(s1)}$$

The background absorbance was measured from buffer-only controls.

For *B. subtilis* and *C. crecentus* strains, we used a fluorogenic substrate, MUG, which is more sensitive than ONPG[3]. We mixed 100 µL of cell lysate with 400 µL of 1x Z buffer and 50 µL of 2 mg/mL MUG. The samples were incubated at 37°C for 1 to 24 hours. To stop LacZ reaction, 55 µL of the reaction mixture was mixed with 25 µL of 1 M sodium carbonate, and the fluorescence was measured in the microplate reader with a blue filter set (EX 360/40 and EM 460/40). LacZ activity was calculated using the following formula:

$$\text{Activity} = \frac{(\mathrm{F}_{460} - F_{460,background})}{\text{Reaction time} \cdot (\mathrm{OD}_{600} - OD_{600,background})} \quad \text{(s2)}$$

The background fluorescence was measured from buffer-only controls. For *C. crescentus*, $OD_{660}$ was used instead of $OD_{600}$.

Once LacZ activities were plotted over sampling times, the baseline was determined by averaging LacZ activities between t = 0 (time of IPTG addition) and the start of LacZ appearance. This baseline activity was subtracted from each point to obtain the change in LacZ activity from the basal level (e.g., **Fig. 6E**). To calculate translation time (when the first LacZ proteins appear), we plotted the square root of LacZ activity (after baseline subtraction) and performed a linear fit to the initial increase to find the intercept[4] (**Fig. S6B** and **S6F**).

When *lacZ* expression was re-repressed at t = 75 s, LacZ protein levels reached a plateau around t = 7 min (**Fig. S4A**). LacZ protein activity at the plateau was calculated by averaging data points between t = 7 and 10 min, and the difference to the baseline was used as a proxy for total LacZ protein expression from 75-s induction (**Fig. 4D** and **S5**).

**Fluorescence *in situ* hybridization (FISH) microscopy**

We followed a procedure previously published[1]. For hybridization, we used twenty-four single-stranded DNA probes complementary to the first 1-kb regions of *lacZ* mRNA ($Z5_{FISH}$). Each probe was labeled with a single Cy3B at the 5' end. The sequences are provided in **Table S3**.

Phase contrast and fluorescence microscopy were performed on an Eclipse Ti-2 microscope (Nikon) equipped with a Sola SE II 365 light engine (Lumencor), a phase-contrast objective Plan Apochromat 100×/1.45 NA (Nikon), and an Orca-R2 CCD camera (Hamamatsu Photonics). All images were acquired using the Nikon Elements software (Nikon).

For analysis, cell outlines were obtained from phase contrast images using the cell detection module of the open-source image analysis software Oufti[5]. FISH fluorescent spots were identified using the open-source tracking software u-track[6]. To determine the fluorescent spot location and intensity, we employed the Gaussian mixture-model fitting method in the single-particle detection module of u-track. The Gaussian standard deviation of a point spread function was calculated based on 0.21*emission wavelength of Cy3 (570 nm)/numerical aperture of the objective lens (1.45). We used an alpha value of 0.015 in the hypothesis test to pick up local maxima significantly brighter than the background. For the Gaussian fitting of local maxima, we applied the default alpha value of 0.05 to test whether the amplitude of the fitted Gaussian and the distance between two fitted Gaussian centers are significantly different from zero. The same set of parameter values was used to analyze data from different time points, days of experiments, and strains.

To combine the analysis results of Oufti and u-track, custom-written MATLAB code was used (see **Software availability**). In the code, fluorescent spots enclosed by cell outlines were assigned to corresponding cells. This allows for analyzing statistics, such as the number of spots per cell (**Fig. 5E**). To analyze the subcellular localization of mRNA, the positions of fluorescent spots along the long and short axes of the corresponding cell were calculated and normalized to cell length and width, respectively. For statistics, the distribution of the normalized positions was plotted as a 2D histogram (**Fig. 3B**-**3C**, **3F**-**3G**, and **5C**-**5D**). For the 2D histograms, the same color bar scheme was used for all strains.

**Mathematical model for the temporal changes in mRNA expression levels during induction and re-repression experiments**

The temporal change in the mRNA level, as shown in **Figure 1C** and **S1A**, was obtained by solving ordinary differential equations (ODE) of chemical rates for nascent and released mRNAs. We denote the time that the first RNAP passes Z5 and Z3 probe regions as $T_{5'}$ and $T_{3'}$, respectively. We denote the time that the last RNAP passes Z5 and Z3 probe regions as $t_{5'}$ and $t_{3'}$, respectively (**Fig. 1B**-**1C**). Even though Z3 probe is close to the end of the gene, we introduce two new variables to indicate the time that the first and last RNAPs reach the end of the gene (intrinsic terminator site) as $T_{end}$ and $t_{end}$. For example, for the wild-type strain (SK98), we estimate that $T_{end}$ is about 15 s later than $T_{3'}$ because Z3 probe is 222 bp before the end of the gene and RNAP speed is about 13.5 nt/s based on $T_{5'}$ and $T_{3'}$.

In our re-repression experiments, we added glucose (or rifampicin) soon after induction, such that the reference time points are in the following order:

$$T_{5'} < t_{5'} < T_{3'} < T_{end} < t_{3'} < t_{end}$$

Then, ODEs for nascent and released mRNAs can be written at different time windows. Below $k_i$ is transcription initiation rate at the promoter, and $k_{d1}$ and $k_{d2}$ are the degradation rate of nascent and released mRNAs.

| For Z5 | Nascent mRNA, $m_n$ | Released mRNA, $m_r$ |
|---|---|---|
| $T_{5'} \leq t \leq t_{5'}$ | $\frac{dm_n(t)}{dt} = k_i - k_{d1}m_n(t)$ …(s3) | $\frac{dm_r(t)}{dt} = 0$ …(s4) |
| $t_{5'} \leq t \leq T_{end}$ (Time window i) | $\frac{dm_n(t)}{dt} = -k_{d1}m_n(t)$ …(s5) | $\frac{dm_r(t)}{dt} = 0$ …(s6) |
| $T_{end} \leq t \leq t_{end}$ (Time window ii) | $\frac{dm_n(t)}{dt} = -k_i e^{-d_{m1}(T_{end}-T_{5'})} - k_{d1}m_n(t)$ …(s7) | $\frac{dm_r(t)}{dt} = k_i e^{-d_{m1}(T_{end}-T_{5'})} - k_{d2}m_r(t)$ …(s8) |
| $t_{end} \leq t$ (Time window iii) | $\frac{dm_n(t)}{dt} = 0$ …(s9) | $\frac{dm_r(t)}{dt} = -k_{d2}m_r(t)$ …(s10) |

Furthermore, we can write ODEs for Z3 signal. We assumed its co-transcriptional and post-transcriptional degradation rates are the same as those of Z5.

| For Z3 | Nascent mRNA, $m_n$ | Released mRNA, $m_r$ |
|---|---|---|
| $T_{3'} \le t \le T_{end}$ | $\frac{dm_n(t)}{dt} = k_i - k_{d1}m_n(t)$ …(s11) | $\frac{dm_r(t)}{dt} = 0$ …(s12) |
| $T_{end} \le t \le t_{3'}$ | $\frac{dm_n(t)}{dt} = 0$ …(s13) | $\frac{dm_r(t)}{dt} = k_i e^{-k_{d1}(T_{end}-T_{3'})} - k_{d2}m_r(t)$ …(s14) |
| $t_{3'} \le t \le t_{end}$ | $\frac{dm_n(t)}{dt} = -k_i e^{-k_{d1}(T_{end}-T_{3'})} - k_{d1}m_n(t)$ …(s15) | $\frac{dm_r(t)}{dt} = k_i e^{-k_{d1}(T_{end}-T_{3'})} - k_{d2}m_r(t)$ …(s16) |
| $t_{end} \le t$ | $\frac{dm_n(t)}{dt} = 0$ …(s17) | $\frac{dm_r(t)}{dt} = -k_{d2}m_r(t)$ …(s18) |

The 4th order Runge-Kutta method was used to obtain numerical solution of these ODEs. To obtain the initial condition, we used the following two equations based on transcription initiation rate at the repressed state ($k_{basal}$).

$$m_n(t=0) = m_n(t=T_{5'}) = (k_{basal}/k_{d1})[1 - e^{-k_{d1}(T_{end}-T_{5'})}] \quad \text{(s19)}$$

$$m_r(t=0) = m_r(t=T_{end}) = (k_{basal}/k_{d2})e^{-k_{d1}(T_{end}-T_{5'})} \quad \text{(s20)}$$

For results plotted in **Figure 1C** and **S1A** (case 1), following parameter values were used:

$$k_{basal} = 0.0064\, min^{-1}, k_i = 1.06\, min^{-1}, k_{d1} = 0.18\, min^{-1}, k_{d2} = 0.42\, min^{-1}$$

For case 2 and case 3 in **Figure S1A**, only $k_{d1}$ was changed to 0 and 0.42 $min^{-1}$, respectively. For Z3 shown in **Figure 1C**, we used:

$$k_{basal} = 0.0099\, min^{-1}, k_i = 0.80\, min^{-1}, k_{d1} = 0.18\, min^{-1}, k_{d2} = 0.42\, min^{-1}$$

**Comparison of translation efficiency of RBS mutants to genome-wide protein-mRNA ratio**

Using *E. coli* YFP library, Taniguchi et al. quantified the number of proteins at the single-molecule and single-cell levels[7]. The number of mRNAs were also counted for a subset of genes at the single-molecule and single-cell levels using FISH, and they were in a good correlation with mRNA levels measured by RNA seq, which covered all expressed mRNAs (as shown in Fig. 3C or Fig. S22 in the paper[7]). In the supplementary data, they provided protein-mRNA ratio for 585 genes, which is a good approximation for the number of proteins made per transcript, or translation efficiency[8], given that their protein numbers were the absolute copy numbers per cell and mRNA numbers (from RNA seq) can be considered as absolute copy numbers per cell as well, based on its good correlation with FISH-based counting data. We plotted this protein-mRNA ratio as a histogram to illustrate variations in translation efficiency across the genome (**Fig. S5H**).

We can also estimate protein-mRNA ratio for *lacZ* in our RBS mutants. For protein counts, we estimated the protein expression level (arbitrary unit) from 75-s induction by Miller assay (see example data in **Fig. 4D** and **S4A**). For the mRNA level, we used the steady-state level of Z3 from no-glucose experiment (arbitrary unit). We used Z3 instead of Z5, because some Z5 signal may include prematurely terminated transcripts, which do not produce full LacZ proteins. After taking the ratio between these two observables, we multiplied a constant to make the protein-mRNA ratio for SK98, the original RBS of *lacZ* is 20. This number came from another single-molecule counting study showing that one *lacZ* mRNA produces about 20 monomers of LacZ protein[9]. These protein-mRNA ratio estimates for various RBS mutants are shown in **Figure S5G**-**S5H**.

**Calculation of the probability of premature termination during transcription**

We investigated how to calculate the probability of premature transcription termination using the steady-state levels of Z5 and Z3. To do so, we first derived generalizable formulae for the steady-state levels of 5' and 3' mRNA. The following derivation assumes that co-transcriptional mRNA degradation is negligible, and hence it is suitable for *E. coli* and *B. subtilis* cases.

We will denote 5' mRNA signal (e.g. Z5) as $N_5$ and 3' mRNA signal (e.g. Z3) as $N_3$. $N_5$ is the sum of three different mRNA species: (a) nascent mRNAs ($N_{5N}$), (b) prematurely terminated mRNAs ($N_{5PT}$), and (c) released mRNAs after full-length transcription ($N_{5R}$). Similarly, $N_3$ can be written as the sum of two different species: (a) nondegraded, released mRNAs ($N_{3R}$) and (b) released mRNAs with the 5' part degraded ($N_{3D}$).

$$N_5 = N_{5N} + N_{5PT} + N_{5R} \tag{s21}$$

$$N_3 = N_{3R} + N_{3D} \tag{s22}$$

Note that we ignored $N_3$ coming from nascent mRNAs because Z3 probes are very near the end of the gene. By definition, $N_{3R}$ is equal to $N_{5R}$. $N_{3D}$ is to reflect mRNA degradation following 5' → 3' direction[10-12]. Such directional decay was needed to explain equal steady-state levels of Z5 and Z3 in the case of 0% premature termination; e.g., native *lacZ* strain (SK98; **Fig. 5F**) and weak RBS strain treated with BCM (SK421+BCM; **Fig. 5H**). If we rather assume that 5' and 3' regions of the released *lacZ* mRNA decay independently from each other (at the rate of $k_{d2}$ for each), the steady-state level of Z5 should be higher than that of Z3 in the case of 0% premature transcription termination. This is because nascent mRNAs contribute to Z5 but not to Z3 while released mRNAs contribute to Z5 and Z3 levels equally.

The rate of change of $N_{5R}, N_{3R}$ and $N_{5PT}$ can be expressed as:

$$\frac{dN_{5PT}}{dt} = k_i \cdot PT - N_{5PT} \cdot k_{dPT} \tag{s22}$$

$$\frac{dN_{5R}}{dt} = k_i \cdot (1 - PT) - N_{5R} \cdot k_{d2} \tag{s23}$$

$$\frac{dN_{3R}}{dt} = k_i \cdot (1 - PT) - N_{3R} \cdot k_{d2} \tag{s24}$$

where $k_i$ is the transcription initiation rate at the promoter, *PT* is the fraction of RNAPs that are prematurely terminated, (1-*PT*) is the fraction of RNAPs that complete full-length transcription, $k_{dPT}$ is the degradation rate of prematurely terminated mRNAs, and $k_{d2}$ is the post-transcriptional degradation rate of fully transcribed mRNAs. The first terms in equation (s22)-(s24) are related to the introduction of new mRNAs into the category. If nascent mRNAs are degraded co-transcriptionally (with rate $k_{d1*}$), the first terms should be multiplied by an exponential factor, $\exp(-k_{d1*}\Delta t)$, where $\Delta t$ is the time duration that the mRNA stay nascent (e.g., for $N_{5PT}$, it is the time for an RNAP to translocate from Z5 probe region to the site of premature termination). For the sake of simplicity, we assumed this factor is 1, and it is justified by our finding that co-transcriptional degradation is more than 10 time slower than post-transcriptional degradation for genes encoding cytoplasmic proteins. From equation (s22)-(s24), we get the steady-state solution for $N_{5PT}$, $N_{5R}$, and $N_{3R}$.

$$N_{5PT} = \frac{k_i}{k_{dPT}} \cdot PT \tag{s25}$$

$$N_{5R} = N_{3R} = \frac{k_i}{k_{d2}}(1 - PT) \tag{s26}$$

The steady-state level of $N_{5N}$ is equal to the number of RNAPs on the gene because co-transcriptional mRNA degradation is close to zero (**Fig. 1D**). First, we find the time ($t_{RNAP}$) that one RNAP takes to travel the distance between 5'-end and 3'-end probes using the distance between the probes ($L$) and RNAP speed ($v_{RNAP}$). Multiplying the RNAP loading rate ($k_i$) by the time $t_{RNAP}$ gives the number of RNAPs on the DNA in the absence of premature transcription termination.

$$t_{RNAP} = \frac{L}{v_{RNAP}} \tag{s27}$$

$$N_{5N} = \frac{L}{v_{rnap}} k_i \tag{s28}$$

In the presence of premature transcription termination, $N_{5N}$ will be different. For simplicity, we assume that premature termination takes place in the middle of the gene, at *L*/2. Effectively, this scenario gives the same result as having sequence-independent premature termination (occurring anywhere with equal probability), which was observed when transcription became uncoupled to translation in the presence of translation inhibiting drug, fusidic acid[13]. Then, we can break down the nascent mRNA population into two populations: those occurring before or after the *L*/2 position. $N_{5N}$ can be written as the sum of the two as:

$$N_{5N} = \frac{L}{2v_{rnap}} k_i + \frac{L}{2v_{rnap}} k_i(1 - PT) = \frac{L}{2v_{rnap}} k_i(2 - PT) \tag{s29}$$

To calculate the steady-state level of $N_{3D}$, we denote the amount of time it takes for the degradation to reach Z3 after Z5 is degraded as $t_D$. $t_D$ is multiplied by the rate of mRNAs entering degradation, given by the decay term ($N_{3R} \cdot k_{d2}$) in equation (s24). Then, the steady-state level of $N_{3D}$ can be expressed as:

$$N_{3D} = t_D \cdot N_{3R} \cdot k_{d2} = t_D \cdot k_i(1 - PT) \tag{s30}$$

Combining all the components for $N_5$ and $N_3$, we get the following equations for the steady state:

$$N_5 = N_{5N} + N_{5PT} + N_{5R} = \frac{L}{2v_{rnap}} k_i(2 - PT) + \frac{k_i}{k_{dPT}} \cdot PT + \frac{k_i}{k_{d2}}(1 - PT) \quad \text{(s31)}$$

$$N_3 = N_{3R} + N_{3D} = \frac{k_i}{k_{d2}}(1 - PT) + t_D \cdot k_i(1 - PT) \quad \text{(s32)}$$

When there is no premature transcription termination ($PT$ =0, $N_{5PT} = 0$), we find that $N_{3D}$ should be equal to $N_{5N}$ in order to get $N_5 = N_3$ at the steady state. This suggests, $t_D = t_{RNAP}$, or the 5'→3' directional degradation follows the elongation rate of RNAP.

Finally, we examine how the steady-state $N_5$ and $N_3$ ratio (which we can obtain from qRT-PCR data, e.g., **Fig. 5F**-**5H**) is related to the probability of premature transcription termination, *PT*. Using equations (s31) and (s32),

$$\frac{N_5 - N_3}{N_5} = \frac{\frac{L}{2v_{rnap}} \cdot PT + \frac{1}{k_{dPT}} \cdot PT}{\frac{L}{2v_{rnap}}(2 - PT) + \frac{1}{k_{dPT}} \cdot PT + \frac{1}{k_{d2}} \cdot (1 - PT)} \quad \text{(s33)}$$

From experiments, we obtained $t_{RNAP} = L/v_{RNAP}$ as 90 s (**Fig. 5F**) and $k_{d2}$ as 0.4 min$^{-1}$ (**Fig. 5K**). Then, the right-hand side depends on *PT* and $k_{dPT}$ (**Fig. S5A**).

Assuming an initial value for $k_{dPT}$, we converted the steady-state levels of $N_5$ and $N_3$ into *PT* using equation (s33). By fitting $k_{d1}$ vs these new *PT* values using $k_{d1} = k_{d1*}(1 - PT) + k_{dPT} \cdot PT$ (equation (1) in the main text), we can get $k_{d1*}$ and $k_{dPT}$. Then, we repeated this process iteratively until the $k_{dPT}$ values converged. We obtained $k_{dPT}$ = 0.799 ± 0.0587 min$^{-1}$ and $k_{d1*}$= 0.0248 ± 0.0372 min$^{-1}$, as well as *PT* values for each strain (**Fig. 5J**; see **Software availability**).

Error bars in *PT* values in **Figure 5J** were calculated using the standard deviation in the ratio of $N_5$ and $N_3$ levels at the steady state. The uncertainties in $k_{dPT}$ and $k_{d1*}$ are one standard deviation from least-square fitting (Igor Pro, Wavemetrics).

If premature transcription termination takes place (a) right after Z5 or (b) right before Z3, equation (s29) will be modified as:

$$N_{5N} = \frac{L}{v_{rnap}} k_i(1 - PT) \quad \text{(s29a)}$$

$$N_{5N} = \frac{L}{v_{rnap}} k_i \quad \text{(s29b)}$$

These are two extreme cases in contrast to the scenario we considered in the equation (s29). If we use one of these equations in place of (s29), we get $k_{dPT}$ = 0.717 min$^{-1}$ and $k_{d1*}$= 0.0203 min$^{-1}$ (in case premature termination happens right after Z5, with rate *PT*) or $k_{dPT}$ = 0.881 min$^{-1}$ and $k_{d1*}$= 0.0290 min$^{-1}$ (in case premature termination happens right before Z3, with rate *PT*). These results are similar to our original conclusion using equation (s29): $k_{dPT}$ = 0.799 min$^{-1}$ and $k_{d1*}$= 0.0248 min$^{-1}$, suggesting that the exact location of premature transcription termination is not critical to our conclusion.

**Calculation of the probability of premature transcription termination in *C. crescentus***

To accurately calculate the probability of premature transcription termination during transcription in *C. crescentus* based on the steady-state levels of Z5 and Z3, we need to modify the equations written for *E. coli*. Because in *C. crescentus* RNase E is localized in the cytoplasm, co-transcriptional mRNA degradation may not be negligible. We observed that in *C. crescentus*, Z5 reaches a steady state before Z3 appears from the baseline (**Fig. 6F** and **S6J**). This indicates that synthesis and degradation of Z5 become balanced even before the first Z3 is synthesized (when all Z5 are supposed to be nascent), indicating significant co-transcriptional mRNA degradation and premature transcription termination. Based on this result, we assume that there is no full-length mRNA released with 5' present. Incorporating these ideas, we write $N_5$ as the sum of two different mRNA species: (a) nascent mRNAs ($N_{5N}$) and (b) prematurely terminated mRNAs ($N_{5PT}$). $N_3$ is from one species, released mRNAs which have the 5' part degraded ($N_{3R}$).

$$N_5 = N_{5N} + N_{5PT} \quad \text{(s34)}$$

$$N_3 = N_{3R} \quad \text{(s35)}$$

We ignored $N_3$ coming from nascent mRNAs because Z3 probes are very near the end of the gene.

The rate of change of $N_{5PT}$ and $N_{3R}$ can be expressed as:

$$\frac{dN_{5PT}}{dt} = k_i \cdot PT \cdot e^{-k_{d1*}\tau} - N_{5PT} \cdot k_{dPT} \quad \text{(s36)}$$

$$\frac{dN_{3R}}{dt} = k_i \cdot (1 - PT) - N_{3R} \cdot k_{d2} \quad \text{(s37)}$$

where $k_i$ is the transcription initiation rate at the promoter, *PT* is the fraction of RNAPs that are prematurely terminated, (1-*PT*) is the fraction of RNAPs that complete full-length transcription, $k_{d1*}$ is the co-transcriptional degradation rate of nascent mRNA (5' end), $k_{dPT}$ is the degradation rate of prematurely terminated mRNAs (5' end), and $k_{d2}$ is the post-transcriptional degradation rate of released mRNA (3' end or Z3). The first term in equation (s36) has an exponential factor, $e^{-k_{d1*}\tau}$, where $\tau$ is the time duration that the mRNA stays nascent. Assuming that the premature termination takes place in the middle of the gene ($x = L/2$), this duration is determined by the RNAP travel time from the Z5 probe region to the site of premature termination.

$$\tau = \frac{L}{2v_{RNAP}} \quad \text{(s38)}$$

At the steady state,

$$N_{5PT} = \frac{k_i}{k_{dPT}} \cdot PT \cdot e^{-k_{d1*}\tau} \quad \text{(s39)}$$

$$N_{3R} = \frac{k_i}{k_{d2}} (1 - PT) \quad \text{(s40)}$$

The number of nascent mRNA with Z5 at the steady state, $N_{5N}$, is related to the density of RNAPs along the DNA length, given by $\frac{k_i}{v_{rRNAP}}$ (the number of RNAPs per unit length). When we incorporate the effect of co-transcriptional degradation reducing $N_{5N}$, the expression for density of $\frac{dN_{5N}}{dx}$ along the DNA length can be written as:

$$\frac{dN_{5N}}{dx} = \begin{cases} \frac{k_i}{v_{RNAP}} e^{-k_{d1*}x/v_{RNAP}} \left(for\ x \leq \frac{L}{2}\right) \\ \frac{k_i}{v_{RNAP}} (1-PT)\, e^{-k_{d1*}x/v_{RNAP}} \left(for\ x \geq \frac{L}{2}\right) \end{cases} \quad \text{(s41)}$$

where $x$ is the base pair position of an RNAP along the DNA ranging from $0 \leq x \leq L$ and can be defined as $x = v_{RNAP} \cdot t$. These expressions can then be integrated along the length of the DNA between the Z5 and Z3 probe positions to obtain the amount of nascent mRNA.

$$N_{5N} = \int_0^{L/2} \frac{k_i}{v_{RNAP}} e^{-k_{d1*}x/v_{RNAP}}\, dx + \int_{L/2}^{L} \frac{k_i}{v_{RNAP}} (1-PT) e^{-k_{d1*}x/v_{RNAP}}\, dx \quad \text{(s42)}$$

$$N_{5N} = \frac{k_i}{k_{d1*}} (1 - e^{-k_{d1*}\tau})[1 + (1-PT)e^{-k_{d1*}\tau}] \quad \text{(s43)}$$

Notably, in the small approximation limit ($k_{d1*} \ll 1$), or in the absence of co-transcriptional mRNA degradation, $e^{-k_{d1*}\tau} \approx 1 - k_{d1*}\tau$, and we can get the equation for $N_{5N}$ used for the *E. coli* case (equation (s29)) from equation (s43).

Putting these together,

$$\frac{N_5 - N_3}{N_5} = \frac{\frac{k_i}{k_{d1*}}(1 - e^{-k_{d1*}\tau})[1 + (1-PT)e^{-k_{d1*}\tau}] + \frac{k_i}{k_{dPT}} \cdot PT \cdot e^{-k_{d1*}\tau} - \frac{k_i}{k_{d2}}(1-PT)}{\frac{k_i}{k_{d1*}}(1 - e^{-k_{d1*}\tau})[1 + (1-PT)e^{-k_{d1*}\tau}] + \frac{k_i}{k_{dPT}} \cdot PT \cdot e^{-k_{d1*}\tau}} \quad \text{(s44)}$$

The steady-state ratio (left-hand side) is measured from the experiment to be about 0.16 ± 0.08 (**Fig. 6F**). $\tau$ is approximated to be 69 s (based on transcription time of 2.3 min; **Fig. 6F**). $k_{d2}$ is about 0.38 min$^{-1}$ (**Fig. S6J**-**6K**). Then, there are three unknowns in this equation (s44): *PT*, $k_{d1*}$, and $k_{dPT}$. These unknown variables are also related to each other for $k_{d1}$, which is measured to be 1.28 min$^{-1}$ (**Fig. 6G**), and the relationship is stated in equation (1) in the main text.

Note that equation (1) of the main text is a good approximation when co-transcriptional degradation is minimal. For *C. crescentus*, this needs to be updated to account for the effect of co-transcriptional degradation on prematurely terminated transcripts. This is because among prematurely terminated mRNAs, only those still have Z5 (not degraded by $k_{d1*}$) will contribute to $k_{d1}$.

$$k_{d1} = k_{d1*}(1-PT) + k_{dPT} \cdot PT \cdot e^{-k_{d1*}\tau} \quad \text{(s45)}$$

When premature transcription termination is blocked by BCM (*PT* = 0), we measured $k_{d1} = k_{d1*} = 0.71 \pm 0.093 \text{ min}^{-1}$ (**Fig. S6I**). Using this value, we analytically solved equation (s44) and (s45) for remaining unknowns: *PT* and $k_{dPT}$ (see **Software availability**). The uncertainties in *PT* and $k_{dPT}$ are estimated by using measurement errors in constant variables used in the two equations, including the steady-state ratio, $\tau$, $k_{d2}$, and $k_{d1*}$.Specifically, each of these parameter values was sampled from a normal distribution defined by its mean and standard deviation acquired from the experiment.

##### Statistical test

Two-sample unpaired Student's *t* tests were performed by using the MATLAB ttest2 function (**Table S7**).

##### Software availability

MATLAB code for data analysis and python code for modeling are available at the publicly accessible site, https://github.com/sjkimlab/Code_Publication.

#### Supplemental Tables

**Table S1. Bacterial strains and plasmids used in this study.**

| **Strain number** | **Genotype** | **Source (reference)** | **Use in this study** |
|---|---|---|---|
| ***E. coli* strains** | | | |
| CJW5615 | MG1655 Δ*lacIZYA*::*kan* | C. Jacobs-Wagner | Cloning |
| CJW5685 | MG1655 *rne*::*rne*ΔMTS-*mcherry* | C. Jacobs-Wagner | Cloning |
| CJW6643 | MG1655 Δ*lacIZYA* | C. Jacobs-Wagner | Cloning |
| JRH471 | MG1655 *rne*(1-529)-*yfp kan* | C. Jacobs-Wagner | Cloning |
| JRH474 | MG1655 *rne*(1-592)-*yfp kan* | C. Jacobs-Wagner | Cloning |
| SK24 | MG1655 *rne*3071 *zce*-726::Tn10 | C. Jacobs-Wagner | Cloning |
| SK52 | MG1655 Δ*araFGH* P13-*araE* | C. Jacobs-Wagner | Cloning |
| SK98 | MG1655 Δ*lacYA* | C. Jacobs-Wagner CJW5461[1] | Cloning, qRT-PCR, Miller assay, FISH |
| SK301 | MG1655 Δ*lacYA* Δ*rhlB*::*kan* | This study | qRT-PCR |
| SK306 | MG1655 Δ*lacYA* Δ*pnp*::*kan* | This study | qRT-PCR |
| SK337 | MG1655 *rne*::*rne*ΔMTS *kan* | This study | Cloning |
| SK339 | MG1655 Δ*lacYA* *rne*::*rne*ΔMTS *kan* | This study | qRT-PCR |
| SK369 | MG1655 Δ*lacYA* *rne*::*rne*(1-529)-*yfp kan* | This study | qRT-PCR |
| SK370 | MG1655 Δ*lacYA* *rne*::*rne*(1-592)-*yfp kan* | This study | qRT-PCR |
| SK390 | MG1655 Δ*lacYA*::$P_{M1}$-*aadA* | This study | qRT-PCR, FISH |
| SK420 | MG1655 Δ(*lacZ* 5’UTR)::weakRBS1 Δ*lacYA* | This study | qRT-PCR, Miller assay |
| SK421 | MG1655 Δ(*lacZ* 5’UTR)::weakRBS2 Δ*lacYA* | This study | Cloning, qRT-PCR, Miller assay |
| SK435 | MG1655 Δ*lacYA*::$P_{M1}$-*lacY*-*venus* | This study | qRT-PCR, FISH |
| SK468 | MG1655 Δ*araFGH* P13-*araE* Δ*lacYA* | This study | Cloning |
| SK472 | MG1655 Δ*araFGH* P13-*araE* *araBAD*::*araB*-*term2* | This study | qRT-PCR |
| SK475 | MG1655 Δ*lacYA*::*kan* | This study | Cloning |
| SK476 | MG1655 Δ*araFGH* P13-*araE* Δ*lacIZYA araBAD*::*lacZ kan* | This study | Cloning |
| SK477 | MG1655 Δ*araFGH* P13-*araE* Δ*lacIZYA araBAD*::*lacZ* | This study | qRT-PCR, Miller assay |

| | | | |
|---|---|---|---|
| SK499 | MG1655 Δ*lacIZYA araBAD*::*lacI-lacZ* | This study | qRT-PCR, Miller assay |
| SK518 | MG1655 Δ*araFGH* P13-*araE* Δ*lacIZYA araBAD*::RBS3-*lacZ* | This study | qRT-PCR, Miller assay |
| SK519 | MG1655 *rne*3071 *zce*-726::Tn10 Δ*lacYA* | This study | qRT-PCR, FISH |
| SK532 | MG1655 Δ*araFGH* P13-*araE* Δ*lacIZYA araBAD*::RBS4-*lacZ* | This study | qRT-PCR, Miller assay |
| SK562 | MG1655 Δ*lacZYA*::*cat* | This study | Cloning |
| SK564 | MG1655 *lacZYA*::*lacY-venus*-term | This study | qRT-PCR |
| SK575 | MG1655 *lacZYA*::*lacY2-lacZ-venus*-term | This study | qRT-PCR, FISH |
| SK591 | MG1655 *rne*3071 *zce*-726::Tn10 Δ(*lacZ* 5'UTR)::weakRBS2 Δ*lacYA* | This study | qRT-PCR, FISH |
| SK613 | MG1655 Δ*araFGH* P13-*araE* Δ*lacIZYA araBAD*::RBS2-*lacZ* | This study | qRT-PCR, Miller assay |
| SX259 | BW25993 *malI*::*malI tetR-eyfp kan lacI*::tetO$_6$-*cat lacI hns*::*hns-mEOS2* | X.S.Xie[14] | Gene-loci imaging |
| SX700 | BW25993 *lacY*::*lacY-venus kan* | X.S. Xie[15] | Cloning |
| ***B. subtilis* strain** | | | |
| GLB503 | 168 Δ*amyE*::pSpankHy-*lacZ* | G.-W. Li[16] | qRT-PCR, Miller assay |
| ***C. crescentus* strain** | | | |
| LS2370 | NA1000 Pxyl::*xylX-lacZ* | L. Shapiro[17] | qRT-PCR, Miller assay |
| **Plasmids** | | | |
| pBAD18kan | | [18] | |
| pCP20 | | [19] | |
| pEXT22 | | [20] | |
| pKD3 | | [19] | |
| pKD4 | | [19] | |
| pKD13 | | [19] | |
| pKD46 | | [19] | |
| pUC19 | | | |
| CJW6650 | pUC19-lacI(O3-)-P$_{M1}$-mgfp(forward)-lacZYA | C. Jacobs-Wagner[1] | |
| SJK1595 | pEXT22-lacI-P$_{M1}$-aadA-term3-lacZ | This study | |
| SJK1758 | pUC19-lacY-venus-term kan | This study | |

| | | |
|---|---|---|
| SJK1761 | pUC19-lacY2-lacZ-venus-term kan | This study |
| SK440 | pBAD18 venus-term2-kan | This study |

##### Construction of plasmids

**SJK1595** plasmid (pEXT22-lacI-$P_{M1}$-aadA-term3-lacZ) was constructed by Gibson assembly of lacI-$P_{M1}$ from CJW6650, *aadA* (spectinomycin resistance gene), and term3-*lacZ* from CJW6650[1]. As a result, three intrinsic terminator sequences (term3) were placed after *aadA*. $P_{M1}$ is a constitutively active promoter[1].

**SJK1758** plasmid (pUC19-lacY-venus-term kan) was constructed by Gibson assembly. *lacY-venus* sequence was amplified from SX700[15], and the kanamycin resistance cassette (*kan*) was amplified from pKD4. The intrinsic transcription terminator sequence (term) after *venus* was from the 3' UTR sequence of *lacA* in the chromosome of MG1655.

**SJK1761** plasmid (pUC19-lacY2-lacZ-venus-term kan) was constructed by Gibson assembly. The first 74 amino acids of LacY, yielding two transmembrane segments[21] (or *lacY*2), were fused to the full-length LacZ (1024 residues), followed by Venus. For intrinsic transcription terminator sequence, the venus gene was followed by 3' UTR sequence of *lacA* in MG1655.

**SK440** plasmid (pBAD venus-term2-kan) was constructed by Gibson assembly of two intrinsic terminators (rrnB T1 and rrnB T2 from pBAD18[18]) and *kan* (pKD4).

##### Construction of strains

**SK301** FRT-*kan*-FRT from pKD13 was amplified with rhlB_KO_F and rhlB_KO_R and integrated into SK98 to delete *rhlB*. These primers were previously used to delete *rhlB* in the Keio collection[22].

**SK306** FRT-*kan*-FRT from pKD13 was amplified with pnp_KO_F and pnp_KO_R and integrated into SK98 to delete *pnp*. These primers were previously used to delete *pnp* in the Keio collection[22].

**SK337** The *kan* cassette in pKD13 was amplified using rne_3UTR_F1A and rne_3UTR_R1. These primers contain the 3' UTR sequence of *rne*. Next, the PCR product was used as a template for next PCR using rne_3UTR_F2A and rne_3UTR_R1 to have an overhang for lambda red recombination into CJW5685 (MG1655 *rne*::*rne*ΔMTS-*mCherry*). The integration occurred at the 3' UTR of *rne* to replace mCherry fusion in CJW5685 with the *kan* cassette.

rne_3UTR_F1A:

TAATAATTAGCTCAAGTAATCAAGCCCTGGTAACTGCCAGGGCTTTTTATTT
CATCTTTGAAATCCGTCGACCTGCAGTTCG

rne_3UTR_R1:

TTGCATTGTTGGTTAGCAAGGATGCCATTCGATGAATTTTAATATGTTGATGT
AGGCTGGAGCTGCTTCG

rne_3UTR_F2A:

CGGCAACACATCATGCCTCTGCCGCTCCTGCGCGTCCGCAACCTGTTGAGT
AATAATTAGCTCAAGTAATCAAGC

**SK339** Phage lysate carrying *rne*::*rne*ΔMTS *kan* (SK337) was transduced into SK98.

**SK369** Phage lysate carrying *rne*::*rne*(1-529)-*yfp kan* (JRH471) was transduced into SK98.

**SK370** Phage lysate carrying *rne*::*rne*(1-592)-*yfp kan* (JRH474) was transduced into SK98.

**SK390** $P_{M1}$-*aadA*-term3 was amplified from SJK1595 plasmid using K074 and K075 and integrated into the downstream of *lacZ* in SK98. Spectinomycin resistance was used for selection.

K074:
CGCTGCGCTTATCAGGCCTACAAGTTCAGCGATCTACATTAGCCGCATCCG
CGATTCTCTTCTGTAAATTGTCG

K075:
TGGGTCAAAGAGGCATGATGCGACGCTTGTTCCTGCGCTTTGTTCATGCCT
TAGTGTATCAACAAGCTGG

**SK420** A synthetic 111-bp DNA (weak RBS1) was integrated into SK98 to change the original RBS sequence of *lacZ* via lambda-red recombination. X-gal blue colony screening was used for selection, and candidate colonies were verified by DNA sequencing.

Weak RBS1:
TTTATGCTTCCGGCTCGTATGTTGTGTGGAATTGTGAGCGGATAACAATTTC
ACACTAAGACCAGCTATGACCATGATTACGGATTCACTGGCCGTCGTTTTAC
AACGTCGTGACTGGGA

**SK421** Same as SK420, using weak RBS2 DNA.

Weak RBS2:
TTTATGCTTCCGGCTCGTATGTTGTGTGGAATTGTGAGCGGATAACAATTTC
ACACGGTTGCCAGCTATGACCATGATTACGGATTCACTGGCCGTCGTTTTAC
AACGTCGTGACTGGGA

**SK435** *lacY-venus kan* was amplified from SX700[15] using K113 and K114 and integrated into SK390 to replace *aadA*. The *kan* cassette was removed by FLP recombination.

K113:
TGTAGACTTTACATCGAATTCGTATTTTGGATGATAACGAGGCGCAAAAAAT
GTACTATTTAAAAAACACAAACTTTTGG

K114:
TGGGTCAAAGAGGCATGATGCGACGCTTGTTCCTGCGCTTTGTTCATGCCT
CGCTGGTGTAGGCTGGAGC

**SK468** First, Δ*lacIZYA*::*kan* (CJW5615) was transduced into SK52 via P1 phage. Then, the *kan* cassette was removed via FLP recombination.

**SK472** term2-*kan* was amplified from SK440 plasmid using K111novenus and K112A and integrated into the chromosome of SK52 (MG1655 Δ*araFGH* P13-*araE*) to replace *araDA*. The *kan* cassette was removed by FLP recombination. We used MG1655 Δ*araFGH* P13-*araE* for homogeneous induction with arabinose in the cell population[23].

K111novenus:

ATCTTCCAACTTCCGCCCCGGCACAGGCTGCCCAGGCCGTTGCGACTCTAT
AACCTGATACAGATTAAATCAGAACG

K112A:

GCTGAAAATCCATCAAAAAACCAGGCTTGAGTATAGCCTGGTTTCGTTTGAT
TGGCTGTGGTTTTATACAGTCAACAGCATATGAATATCCTCCTTAGTTCC

**SK475** The *kan* cassette was amplified from SK440 plasmid using K131 and K132 and integrated into SK98.

K131:

TCTACATTAGCCGCATCCGGCATGAACAAAGCGCAGGAACAAGCGTCGCAT
GTGTAGGCTGGAGCTGCTTCGAAG

K132:

TGCGTTTTGCACCAGTACGTTTTCCGCAGCTGTGGGTCAAAGAGGCATGAC
ATATGAATATCCTCCTTAGTTCC

**SK476** *lacZ-kan* was amplified from SK475 using K125 and K133 and integrated into SK468.

K125:

GCAACTCTCTACTGTTTCTCCATACCCGTTTTTTTGGATGGAGTGAAACGAT
GACCATGATTACGGATTCACTGG

K133:

GCTTGAGTATAGCCTGGTTTCGTTTGATTGGCTGTGGTTTTATACAGTCACA
TGACATATGAATATCCTCC

**SK477** The *kan* cassette was removed from SK476 by FLP recombination to create SK477.

**SK499** *lacI-lacZ-kan* was amplified from SK475 using K136 and K137 and integrated into MG1655 Δ*lacIZYA* (CJW6643, or the *kan*-removed version of CJW5615) to replace *araBAD*. Finally, the *kan* cassette was removed by FLP recombination.

K136:

CGTCACACTTTGCTATGCCATAGCATTTTTATCCATAAGATTAGCGGATCGA
ATCGCAGGCTATTCTGGTGG

K137:

GCTTGAGTATAGCCTGGTTTCGTTTGATTGGCTGTGGTTTTATACAGTCAGA
GGCATGACATATGAATATCC

**SK518** $P_{ara}$-*lacZ-kan* was amplified from SK476 using araBAD_RBS_mut3 and K133 and integrated into the *araB* locus in SK468. The *kan* cassette was removed by FLP recombination.

araBAD_RBS_mut3:

TGACGCTTTTTATCGCAACTCTCTACTGTTTCTCCATACCCGTTTTTTTGAGG
AGCACTGGACGATGACCATGATTACGGATTCACTGG

**SK519** Phage lysate carrying *rne*3071 (SK24) was transduced into SK98.

**SK532** *lacZ-kan* was amplified from SK475 using K146 and K133 to include the original *lacZ* 5' UTR. It was integrated into the *araB* locus in SK468. Finally, the *kan* cassette was removed by FLP recombination.

K146:

AGCGGATCCTACCTGACGCTTTTTATCGCAACTCTCTACTGTTTCTCCATATT
GTGAGCGGATAACAATTTCACACAGG

**SK562** The chloramphenicol resistance (*cat*) cassette was amplified from pKD3 using K162 and K163 and integrated into SK98 to create Δ<u>*lacZYA*</u>.

K162:

ACAGCGCTAGCGGTGATGACGATGACAAGCTGGGAATTGATCCCTTCACCT
CCAAGGGCGAGGAGGATAACC

K163:

CCCACCACCGGAGACGGTGAGAATTTGTCCGCTTACCCAGCTCGCAGCAGT
GAATATCCTCCTTAGTTCC

**SK564** *lacY-venus*-term *kan* was amplified from SJK1758 plasmid using K160 and K132 and integrated into the *lac* locus in MG1655 Δ*lacZYA*::*cat* (SK562). Finally, the *kan* cassette was removed by FLP recombination.

K160:

TATGTTGTGTGGAATTGTGAGCGGATAACAATTTCACACAGGAAACAGCTAT
GTACTATTTAAAAAACACAAACTTTTGG

K132:

TGCGTTTTGCACCAGTACGTTTTCCGCAGCTGTGGGTCAAAGAGGCATGAC
ATATGAATATCCTCCTTAGTTCC

**SK575** Same as SK564, but SJK1761 plasmid was used.

**SK591** Phage lysate carrying *rne*3071 (tet)[24] was transduced into SK421.

**SK613** Same as SK518, but araBAD_RBS_mut2 primer was used.

araBAD_RBS_mut2:

TGACGCTTTTTATCGCAACTCTCTACTGTTTCTCCATACCCGTTTTTTTGTAA
GGCAGGATATTATGACCATGATTACGGATTCACTGG

**Table S2. Oligonucleotides used for RT qPCR (related to Figures 1,2,3,4,5, and 6)**

| Primer | Sequence (5' to 3') |
|---|---|
| lacZ530F | TTTTACGCGCCGGAGAAAAC |
| lacZ530R | AGTCGGTTTATGCAGCAACG |
| lacZ2732F | TTACTGCCGCCTGTTTTGAC |
| lacZ2732R | TGTAGCGGCTGATGTTGAAC |
| lacY80F | TCCCGTTTTTCCCGATTTGG |
| lacY80R_forlacY2 | TTTGCGCAGCCCGAGTTTGTC |
| lacY80R | ACGGCGCAAACATCACTAAC |
| lacY1011F | TACCAGCCAGTTTGAAGTGC |
| lacY1011R | TACATATTGCCCGCCAGTACAG |
| venus585F | GACAACCACTACCTGAGCTA |
| venus585R | CTTGTACAGCTCGTCCATGC |
| araB33F | TGATTCTGTGCGAGCTTTGG |
| araB33R | TGCAAGCACGGTTTTCAGTG |
| araB1536F | AAAAATGGCCAGTGCGGTAG |
| araB1536R | TAGCGGCGATAAAGCTGTTC |

**Table S3. Oligonucleotides used in FISH ($Z5_{FISH}$; related to Figures 3 and 5).**

| Name | Sequence (5’-3’) |
|---|---|
| lacZ1 | GTGAATCCGTAATCATGGTC |
| lacZ2 | TCACGACGTTGTAAAACGAC |
| lacZ3 | ATTAAGTTGGGTAACGCCAG |
| lacZ4 | TATTACGCCAGCTGGCGAAA |
| lacZ5 | ATTCAGGCTGCGCAACTGTT |
| lacZ6 | AAACCAGGCAAAGCGCCATT |
| lacZ7 | AGTATCGGCCTCAGGAAGAT |
| lacZ8 | AACCGTGCATCTGCCAGTTT |
| lacZ9 | TAGGTCACGTTGGTGTAGAT |
| lacZ10 | AATGTGAGCGAGTAACAACC |
| lacZ11 | GTAGCCAGCTTTCATCAACA |
| lacZ12 | AATAATTCGCGTCTGGCCTT |
| lacZ13 | AGATGAAACGCCGAGTTAAC |
| lacZ14 | AATTCAGACGGCAAACGACT |
| lacZ15 | TTTCTCCGGCGCGTAAAAAT |
| lacZ16 | ATCTTCCAGATAACTGCCGT |
| lacZ17 | AACGAGACGTCACGGAAAAT |
| lacZ18 | GCTGATTTGTGTAGTCGGTT |
| lacZ19 | TTAAAGCGAGTGGCAACATG |
| lacZ20 | AACTGTTACCCGTAGGTAGT |
| lacZ21 | ATAATTTCACCGCCGAAAGG |

| | |
|---|---|
| lacZ22 | TTTCGACGTTCAGACGTAGT |
| lacZ23 | ATAGAGATTCGGGATTTCGG |
| lacZ24 | TTCTGCTTCAATCAGCGTGC |

**Table S4. 5'UTR sequence of *lacZ* RBS mutants (related to Figures 4 and 5)**

| Strain | From +1 to start codon |
|---|---|
| SK98 | ATTGTGAGCGGATAACAATTTCACACAGGAAACAGCT**ATG** |
| SK420 | ATTGTGAGCGGATAACAATTTCACACTAAGACCAGCT**ATG** |
| SK421 | ATTGTGAGCGGATAACAATTTCACACGGTTGCCAGCT**ATG** |
| SK477 | ACCCGTTTTTTTGGATGGAGTGAAACG**ATG** |
| SK499 | ATTGTGAGCGGATAACAATTTCACACAGGAAACAGCT**ATG** |
| SK518 | ACCCGTTTTTTTGAGGAGCACTGGACG**ATG** |
| SK532 | ATTGTGAGCGGATAACAATTTCACACAGGAAACAGCT**ATG** |
| SK613 | ACCCGTTTTTTTGTAAGGCAGGATATT**ATG** |

**Table S5. mRNA degradation, premature transcription termination, and protein expression levels of *lacZ* RBS mutants (related to Figure S5)**

| | $k_{d1}$ (Z5) $min^{-1}$ | $k_{d2}$ (Z5) $min^{-1}$ | $k_{d2}$ (Z3) $min^{-1}$ | $\frac{N_5 - N_3}{N_5} * 100$ (%) | Premature termination (%) | LacZ Protein (AU) |
|---|---|---|---|---|---|---|
| SK98 | 0.04±0.060 | 0.43±0.067 | 0.51±0.071 | 0.0±7.7 | 0.0±15 | 24.0±1.40 |
| SK420 | 0.45±0.160 | 0.32±0.091 | 0.51±0.081 | 27±1.6 | 43±2 | 0.2±0.004 |
| SK421 | 0.65±0.171 | 0.52±0.131 | 0.51±0.111 | 56±14 | 72±12 | 0.0±0.00 |
| SK477 | 0.56±0.146 | 0.53±0.009 | 0.52±0.017 | 47±1.1 | 64±1.1 | 0.8±0.14 |
| SK499 | 0.01±0.014 | 0.42±0.034 | 0.54±0.103 | 0±0 | 0.0±0 | 27.3±0.38 |
| SK518 | 0.27±0.086 | 0.39±0.030 | 0.48±0.031 | 34±1.0 | 51±1 | 5.4±0.26 |
| SK532 | 0.04±0.038 | 0.44±0.032 | 0.48±0.037 | 0±0 | 0.0±0 | 47.8±2.01 |
| SK613 | 0.03±0.043 | 0.40±0.034 | 0.48±0.055 | 1.4±1.4 | 2.8±2.7 | 71.6±6.68 |

Error bars represent the standard deviation from three or more biological replicates for $k_d$ values and two biological replicates for LacZ protein. For premature transcription termination, standard deviations in the ratio of steady-state levels of Z5 and Z3 (from two biological replicates) were used to estimate the error range.

**Table S6. Number of spots and cells used in FISH analysis (related to Figures 3 and 5)**

| | Time | Total number of $Z_{5FISH}$ spots | Total number of cells with $Z_{5FISH}$ spot(s) |
|---|---|---|---|
| SK390 | 1 min | 12003 | 8885 |
| | 2 min | 5130 | 3629 |
| | 3 min | 10581 | 7056 |
| SK435 | 1 min | 13945 | 9653 |
| | 2 min | 10606 | 7213 |
| | 3 min | 11101 | 7090 |

| | Time | Total number of $Z_{5FISH}$ spots | Total number of cells with $Z_{5FISH}$ spot(s) |
|---|---|---|---|
| SK575 | 1 min | 7978 | 6087 |
| | 2 min | 12679 | 9203 |
| | 3 min | 7831 | 5721 |
| SK98 | 1 min | 2011 | 1374 |
| | 2 min | 564 | 370 |
| | 3 min | 6801 | 4048 |

| | Time | Total number of $Z_{5FISH}$ spots | Total number of cells with $Z_{5FISH}$ spot(s) |
|---|---|---|---|
| SK519 | 30 s | 29990 | 20311 |
| | 60 s | 39694 | 26197 |
| | 90 s | 48056 | 30800 |
| SK591 | 30 s | 41750 | 26260 |
| | 60 s | 41901 | 24363 |
| | 90 s | 54182 | 30840 |

**Table S7. *P*-values determined by two-tailed Student's *t*-test (related to Figures 2, 3, 4, and 5)**

| Relevant figure | Alternative hypothesis | *P* value |
|---|---|---|
| Figure 2B | $k_{d1}$ of *lacZ* mRNA in WT is smaller than that in RNase E ΔMTS. | 0.0018 |
| Figure 2B | $k_{d2}$ of *lacZ* mRNA in WT is smaller than that in RNase E ΔMTS. | 0.037 |
| Figure 2B | $k_{d1}$ of *lacZ* mRNA in RNase E(1-592) is smaller than that in RNase E(1-529). | 0.026 |
| Figure 2B | $k_{d2}$ of *lacZ* mRNA in RNase E(1-592) is smaller than that in RNase E(1-529). | 0.015 |
| Figure 3D | $k_{d1}$ of *lacZ* mRNA in SK435 (*lacY*) is larger than that in SK390 (*aadA*). | 0.46 |
| Figure 3D | $k_{d2}$ of *lacZ* mRNA in SK435 (*lacY*) is larger than that in SK390 (*aadA*). | 0.93 |
| Figure 3D | $k_{d1}$ of *lacZ* mRNA in SK435 (*lacY*) is larger than that in SK98 (none). | 0.43 |
| Figure 3D | $k_{d2}$ of *lacZ* mRNA in SK435 (*lacY*) is larger than that in SK98 (none). | 0.97 |
| Figure 3H | $k_{d1}$ of *lacZ* mRNA in SK575 (*lacY2-lacZ-venus*) is larger than that in SK98 (none). | 0.73 |
| Figure 3H | $k_{d2}$ of *lacZ* mRNA in SK575 (*lacY2-lacZ-venus*) is larger than that in SK98 (none). | 0.45 |
| Figure 4C | $k_{d1}$ of *araB* mRNA in SK472 is greater than that of *lacZ* mRNA in SK477. | 0.28 |

| | | |
|---|---|---|
| Figure 4C | $k_{d1}$ of *lacZ* mRNA in SK477 is greater than that in SK98. | 0.0040 |
| Figure 4C | $k_{d1}$ of *lacZ* mRNA in SK499 is greater than that in SK98. | 0.77 |
| Figure 4C | $k_{d1}$ of *lacZ* mRNA in SK613 is greater than that in SK499. | 0.28 |
| Figure 4D | LacZ protein expression in SK499 is greater than that in SK477. | 1.84e-06 |
| Figure 4D | LacZ protein expression in SK613 is greater than that in SK499. | 0.011 |
| Figure 5B | $k_{d1}$ of *lacZ* mRNA in SK421 is greater than that in SK98. | 3.9e-04 |
| Figure 5B | $k_{d2}$ of *lacZ* mRNA in SK421 is greater than that in SK98. | 0.15 |
| Figure S1B | $k_{d2}$ of Z5 is greater than $k_{d1}$ in the *lacZ* only strain (SK98). | 1.2e-05 |
| Figure S1B | $k_{d2}$ of Z3 is greater than that of Z5 in the *lacZ* only strain (SK98). | 0.063 |
| Figure S2A | $k_{d1}$ of *lacZ* mRNA in SK98 is larger at 43.5°C than at 30°C. | 0.018 |
| Figure S2A | $k_{d2}$ of *lacZ* mRNA in SK98 is larger at 43.5°C than at 30°C. | 9.9e-04 |
| Figure S2A | At the nonpermissive temperature, $k_{d1}$ of *lacZ* mRNA in the wild-type RNase E strain (SK98) is larger than that in rne3071 strain (SK519). | 0.020 |
| Figure S2A | At the nonpermissive temperature, $k_{d2}$ of *lacZ* mRNA in the wild-type RNase E strain (SK98) is larger than that in rne3071 strain (SK519). | 4.2e-04 |
| Figure S2C | $k_{d1}$ of *lacZ* mRNA in the wild-type strain (SK98) is greater than that in the Δ*rhlB* strain (SK301). | 0.54 |
| Figure S2C | $k_{d2}$ of *lacZ* mRNA in the wild-type strain (SK98) is greater than that in the Δ*rhlB* strain (SK301). | 0.19 |
| Figure S2C | $k_{d1}$ of *lacZ* mRNA in the wild-type strain (SK98) is greater than that in the Δ*pnp* strain (SK306). | 0.39 |
| Figure S2C | $k_{d2}$ of *lacZ* mRNA in the wild-type strain (SK98) is greater than that in the Δ*pnp* strain (SK306). | 0.27 |
| Figure S2C | $k_{d1}$ of *lacZ* mRNA in the wild-type strain (SK98) is greater than that in the RNase E (1-592) strain (SK370). | 0.34 |
| Figure S2C | $k_{d2}$ of *lacZ* mRNA in the wild-type strain (SK98) is greater than that in the RNase E (1-592) strain (SK370). | 0.0020 |
| Figure S3D | $k_{d1}$ of *lacY2* sequence in the complete *lacY* mRNA (SK564) is different from that in *lacY*2-*lacZ-venus* fusion (SK575). | 0.18 |
| Figure S3D | $k_{d2}$ of *lacY2* sequence in the complete *lacY* mRNA (SK564) is different from that in *lacY*2-*lacZ-venus* fusion (SK575). | 0.17 |

| | | |
|---|---|---|
| Figure S4B | $k_{d1}$ of weak RBS-*lacZ* mRNA (SK421) is larger at 43.5°C than at 30°C. | 0.0023 |
| Figure S4B | $k_{d2}$ of weak RBS-*lacZ* mRNA (SK421) is larger at 43.5°C than at 30°C. | 0.89 |
| Figure S4B | At the nonpermissive temperature, $k_{d1}$ of weak-*lacZ* mRNA in the wild-type RNase E strain (SK421) is larger than that in rne3071 strain (SK591). | 5.13e-06 |
| Figure S4B | At the nonpermissive temperature, $k_{d2}$ of weak-*lacZ* mRNA in the wild-type RNase E strain (SK421) is larger than that in rne3071 strain (SK591). | 0.0014 |
| Figure S4B | At the nonpermissive temperature, $k_{d1}$ of weak-*lacZ* mRNA in rne3071 (SK591) is higher than that of native RBS-lacZ mRNA in rne3971 (SK519). | 0.61 |
| Figure S4B | At the nonpermissive temperature, $k_{d2}$ of weak-*lacZ* mRNA in rne3071 (SK591) is higher than that of native RBS-lacZ mRNA in rne3971 (SK519). | 0.068 |
| Figure S6D | $k_{d2}$ of Z5 is larger than $k_{d1}$ in *B. subtilis* (GLB503). | 0.0029 |
| Figure S6D | $k_{d2}$ of Z3 is larger than that of Z5 in *B. subtilis* (GLB503). | 0.63 |
| Figure S6K | For Z5, $k_{d1}$ (measured from adding rifampicin at t = 50 s) is larger than $k_d$ (measured from adding rifampicin at t = 315 s) in *C. crescentus* (LS2370). | 0.28 |
| Figure S6K | $k_d$ of Z5 is larger than $k_{d2}$ of Z3 in *C. crescentus* (LS2370). | 0.020 |

#### Supplemental Figure Legends

**Figure S1** (**A**) Anticipated result of Z5 mRNA levels when *lacZ* expression is induced at t = 0 and re-repressed at t = 75 s. Case 1 represents when co-transcriptional and post-transcriptional degradation take place at distinct rates of $k_{d1}$ = 0.18 min$^{-1}$ and $k_{d2}$ = 0.42 min$^{-1}$. Case 2 represents when co-transcriptional degradation does not take place, i.e., $k_{d1}$ = 0 min$^{-1}$ and $k_{d2}$ = 0.42 min$^{-1}$. Case 3 represents when co-transcriptional degradation takes place at the same rate of post-transcriptional degradation, i.e., $k_{d1} = k_{d2}$ = 0.42 min$^{-1}$. Transcription rate of $k_i$ = 1 min$^{-1}$ was used for the calculation. For transcription elongation, we used $T_{5'}$ = 28 s, $T_{3'}$ = 210 s, $t_{5'}$ = 129 s, $t_{3'}$ = 329 s to match the experimental data shown in **Figure 1D**. (**B**) mRNA degradation rates measured from *lacZ* pulse-induction assay shown in **Figure 1D**. *** denotes p<0.001, and ns indicates a statistically non-significant difference from two-sample t test. Error bars represent the standard deviation from five replicates.

**Figure S2** Effect of RNase E and the RNA degradosome components in *lacZ* mRNA degradation. (**A**) *lacZ* mRNA degradation when RNase E is inactivated. WT strain is SK98. *rne3071* strain is SK519. In all cases, cells were grown at 30°C until they reached OD = 0.2. Only when testing the nonpermissive temperature, cells were shifted to 43.5°C 10 min before induction. Transcription of *lacZ* was induced with 0.2 mM IPTG and re-repressed with 500 mM glucose at t = 75 s (in 30°C experiment), t = 30 s (in 43.5°C experiment of WT *rne*), or t = 50 s (in 43.5°C experiment of *rne3071*). *** and * denote p<0.001 and p<0.05, respectively (two-sample t test). (**B**) Linear representation of RNase E mutants used in **Figure 2**. Numbers are amino acid residues. (**C**) *lacZ* mRNA degradation when one of the RNA degradosome component genes, *rhlB* or *pnp*, is deleted (SK301 and SK306, respectively). WT (SK98) and RNase E (1-592) (SK370) are plotted as a comparison. In all cases, transcription of *lacZ* was induced with 0.2 mM IPTG and re-repressed with 500 mM glucose at t = 75 s. Two-sample t test was performed in comparison to WT (SK98). ** denotes p<0.01, and ns indicates a statistically nonsignificant difference. In panel A and C, error bars represent the standard deviation from three replicates.

**Figure S3** Transertion effect on mRNA degradation. (**A**) Number of $Z5_{FISH}$ fluorescent spots detected per cell at different time points during experiments shown in **Figure 3B-3C**. Error bars represent the standard error from bootstrapping. In each histogram, over 5,000 cells were analyzed. (**B**) Live-cell fluorescence image of LacY2-LacZ-Venus fusion protein in SK575 after induction with 0.2 mM IPTG for 15 min. Scale bar = 1 µm. (**C**) Number of $Z5_{FISH}$ fluorescent spots detected per cell at different time points during experiments shown in **Figure 3F**-**3G**. Error bars represent the standard error from bootstrapping. In each histogram, over 2,000 cells were analyzed, except for SK98 at t = 2 min (370 cells). (**D**) Degradation rate of 5' *lacY* sequence in SK575 (*lacY*2-*lacZ*-*venus*) and SK564 (*lacY*-*venus*), measured from inducing transcription with 0.2 mM IPTG and re-repressing with 500 mM glucose at 75 s (SK575) or 50 s (SK564). qRT-PCR was done using primers amplifying 80-222 nt region (SK575) or 80-268 nt (SK564) of *lacY* sequence. Error bars represent the standard deviation from three replicates. ns indicates a statistically nonsignificant difference (two-sample t test). (**E**) 5' and 3' mRNA level changes after pulse induction of *lacY-venus* under $P_{lac}$ (SK564). 0.2 mM IPTG was added at t = 0, and 500 mM glucose was added at t = 50 s to block further transcription initiation. The 5' sequence was probed by primers amplifying 5' *lacY* sequence (80-268 nt), and the 3' sequence was probed by primers amplifying 585-711 nt region of the *venus* sequence. Blue and yellow areas are where $k_{d1}$ and $k_{d2}$ of 5' *lacY* mRNA were calculated. Error bars represent the standard deviation from three replicates. (**F**) 5' and 3' *lacY* mRNA level changes after induction of *lacY-venus* under $P_{lac}$ (SK564) with 0.2 mM IPTG. 5' and 3' sequences were probed by primers amplifying 80-268 nt and 1011-1118 nt region of the *lacY*, respectively. Error bars represent the standard deviation from two replicates.

**Figure S4** Gene expression difference between native and weak RBS of *lacZ*. (**A**) LacZ protein expression assayed by Miller assay in native and weak RBS *lacZ*. Transcription of *lacZ* was induced with 0.2 mM IPTG at t = 0 and re-repressed with 500 mM glucose at t = 75 s. Error bars represent the standard deviation from two replicates. (**B**) The effect of RNase E on *lacZ* mRNA degradation rate in weak RBS. Cells having the weak RBS sequence for *lacZ* with WT *rne* (SK421) or with *rne*3071 (SK591) were grown at 30°C until they reached OD = 0.2. When non-permissive temperatures were tested, the cell culture was moved to 43.5°C 10 min before induction. Transcription of *lacZ* was induced with 0.2 mM IPTG at t = 0 and repressed with 500 mM glucose at t = 75 s (30°C experiment), t = 30 s (43.5°C experiment for SK421), or t = 50 s (43.5°C experiment for SK591). Native RBS with *rne*3071 (SK519) result was plotted as a comparison. Error bars represent the standard deviation from three or more replicates. *** denotes $p<0.001$, ** denotes $p<0.01$, and ns indicates a statistically nonsignificant difference (two-sample t test). (**C**) Z5 and Z3 mRNA levels in SK591 during the time-course experiment at the nonpermissive temperature, as described in panel B. Blue and yellow areas are where $k_{d1}$ and $k_{d2}$ of Z5 were calculated. Error bars represent the standard deviation of three replicates. (**D**-**E**) Statistics of fluorescent $Z5_{FISH}$ spots at each time point after induction with 0.2 mM IPTG at t = 0 and re-repression with 500 mM glucose at t = 50 s, as described in **Figure 5C**-**5D**. The spots used in the 2D histogram in **Figure 5C**-**5D** were used for the intensity analysis. Error bars represent the standard error from bootstrapping. (**F**) Comparison of the number of *lacZ* gene loci and $Z5_{FISH}$ spots at t = 60 s during the pulse-induction experiment described in **Figure 5C**-**5E**. The gene loci was imaged by TetR-YFP bound to the $tetO_6$ array positioned upstream of *lacZ* in strain SX259[14]. Cells were grown in M9 glycerol at 30°C until OD = 0.2, and shifted to 43.5C for 10 min, similar to the strains used for $Z5_{FISH}$ imaging. Error bars represent the standard error from bootstrapping.

**Figure S5** Relationship between *lacZ* mRNA degradation rates and LacZ protein expression levels in various RBS-*lacZ* mRNA constructs. See **Table S4** for the list of strains used and **Table S5** for data values. *lacZ* mRNA degradation rate was measured from Z5 ($k_{d1}$ and $k_{d2}$) and from Z3 ($k_{d2}$) for each strain from re-repressing transcription with 500 mM glucose at t = 75 s. LacZ protein expression is measured from Miller assay using samples acquired from the same re-repression experiment. A typical Miller assay result is shown in **Figure S4A**. The plateau level (maximum LacZ protein made from the pulsed induction) after the background subtraction is used as a proxy for the protein expression level. (**A**) Theoretical relationship between the probability of premature transcription termination (*PT*), the degradation rate of prematurely released mRNAs ($k_{dPT}$), and the ratio of steady-state levels of Z5 and Z3 ($1-N_3/N_5$). Equation (s33) was used with $k_{d2} = 0.4$ min$^{-1}$ and $t_{RNAP} = 90$ s. Dotted grey line is a visual guide for the case where *PT* equals to $1-N_3/N_5$ (from steady state). (**B**) Relationship between LacZ protein expression level and $k_{d1}$ of Z5. (**C**) Relationship between LacZ protein expression level and $k_{d2}$ of Z5. (**D**) Relationship between LacZ protein expression level and $k_{d2}$ of Z3. (**E**) Relationship between LacZ protein expression level and the probability of premature transcription termination. (**F**) Relationship between post-transcriptional mRNA degradation rates for Z5 and Z3. Dotted line indicates when two rates are equal. (**G**) Relationship between protein-to-mRNA ratio (translation initiation strength or translation efficiency) and probability of premature transcription termination among the RBS mutant strains. Blue shaded area covers strains that we observed nonzero premature transcription termination due to low translation efficiency. Error bars were calculated from propagating the errors in protein levels and the steady-state Z3 mRNA levels, both of which were from the standard deviation from two replicates. (**H**) Histogram of protein-to-mRNA ratio of 585 genes measured in *E. coli* YFP library study[7]. Our RBS mutant strains are marked based on their relative difference to the original *lacZ* (SK98). Blue shaded area indicates genes with low translation efficiency, in which we expect some probability of premature transcription termination based on *PT* values in our RBS mutant strains (panel G). Error bars represent the standard deviation from three replicates (for degradation rates) or two replicates (for LacZ protein and *PT*).

**Figure S6** Expression kinetics of *lacZ* in *B. subtilis* and *C. crescentus*. (**A**) Translation kinetics of *lacZ* in *B. subtilis* probed by Miller assay upon induction with 5 mM IPTG at t = 0. MUG was used as a fluorogenic LacZ substrate. Error bars represent the standard deviation from two replicates. (**B**) Schleif plot[4] of the LacZ induction curve of *B. subtilis* shown in panel A. (**C**) Transcription kinetics of *lacZ* in *B. subtilis* probed by qRT-PCR after induction with 5 mM IPTG. This is a zoomed-in version of **Figure 6B** to show the transcription time (dotted arrow). (**D**) *lacZ* mRNA degradation rates measured from Z5 and Z3 decay in *B. subtilis* from a time-course experiment described in **Figure 6C**. Error bars represent the standard deviation from three replicates. ** denotes $p<0.01$, and ns indicates a statistically nonsignificant difference (two-sample t test). (**E**) LacZ protein expression in *B. subtilis* probed by Miller assay when 200 µg/mL rifampicin was added at t = 30 s after induction with 5 mM IPTG. MUG was used as a sensitive LacZ substrate. Error bars represent the standard deviation from two replicates. (**F**) Schleif plot[4] of the LacZ induction curve of *C. crescentus* shown in **Figure 6E**. (**G**) LacZ protein expression in *C. crescentus* probed by Miller assay when 200 µg/mL rifampicin was added at t = 50 s after induction with 0.3% xylose. MUG was used as a sensitive LacZ substrate. Error bars represent the standard deviation from three replicates. (**H**) *lacZ* mRNA expression upon induction with 0.3 % xylose at t = 0 in the presence of 100 µg/mL BCM, added 5 min before induction. Error bars represent the standard deviations from two replicates. (**I**) *lacZ* mRNA expression upon induction with 0.3 % xylose at t = 0 and re-repression with 200 µg/mL rifampicin at t = 30s in the presence of 100 µg/mL BCM, added 5 min before induction. Error bars represent the standard deviations from three replicates. Blue area indicates where $k_{d1}$, or $k_{d1*}$ of Z5 was calculated without premature transcription termination. (**J**) *lacZ* mRNA level change in *C. crescentus* when 0.3% xylose was added for induction at t = 0 and 200 µg/mL rifampicin was added at t = 315 s. In comparison to the experiment shown in **Figure 6G**, the longer induction helps to develop Z3 signal for its decay measurement. The degradation rate of Z5 and Z3 were measured using an exponential decay fit in the grey box region. Error bars represent the standard deviation from three replicates. (**J**) *lacZ* mRNA degradation rates measured from Z5 and Z3 decay in *C. crescentus*. $k_{d1}$ of Z5 was measured from an experiment shown in **Figure 6G**. $k_d$ of Z5 and $k_{d2}$ of Z3 were measured from panel J. We noted the decay rate of Z5 as $k_d$ because of the ambiguity in assigning them to completely post-transcriptional degradation rate in the experiment shown in panel J. For example, Z5 decays entirely before Z3 starts to decay (around t = 420 s), indicating that Z5 decay around t = 400 s involves co-transcriptional mRNA degradation. Error bars represent the standard deviation from three replicates. * indicates $p<0.05$, and ns indicates a statistically nonsignificant difference (two-sample t test).

### Figure S1

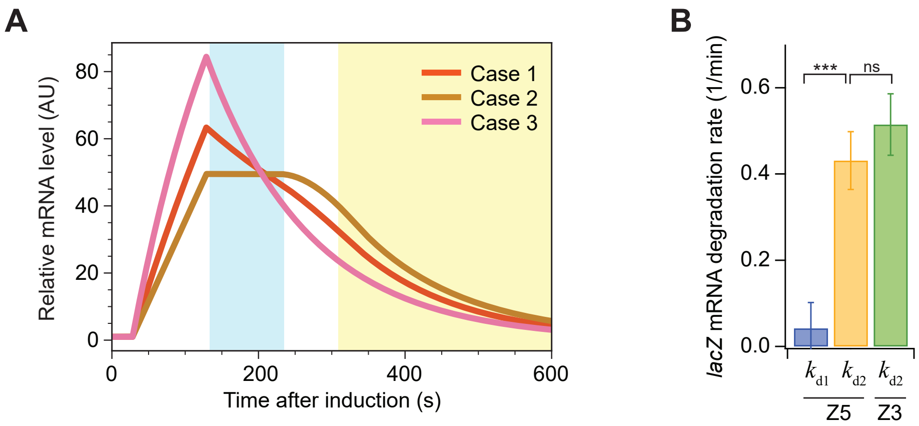
A
Case 1
Case 2
Case 3
Relative mRNA level (AU)
0
20
40
60
80
0
200
400
600
Time after induction (s)
B
lacZ mRNA degradation rate (1/min)
0.0
0.2
0.4
0.6
***
ns
$k_{d1}$
$k_{d2}$
$k_{d2}$
Z5
Z3

### Figure S2

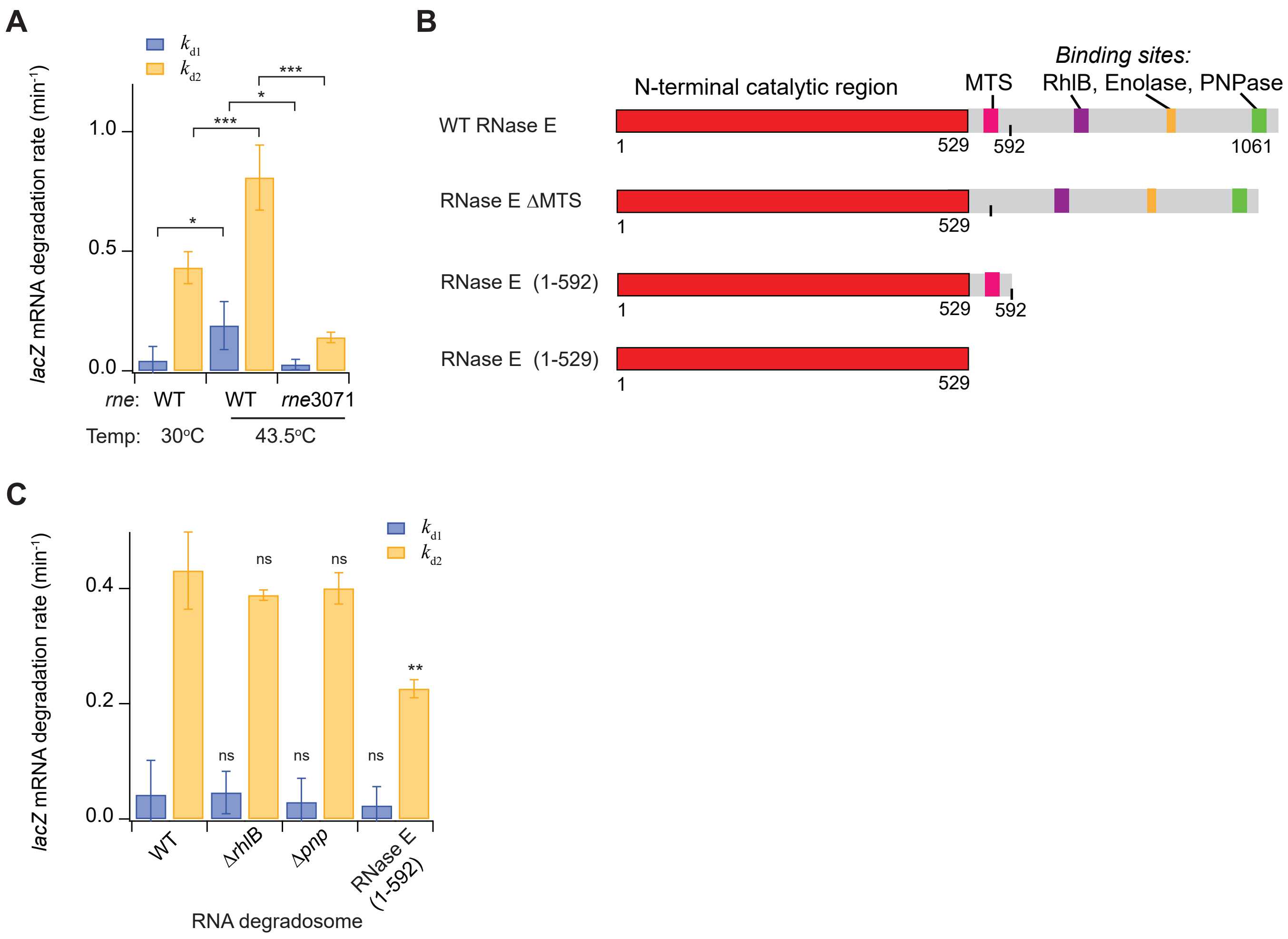
A
B
C
kd1
kd2
lacZ mRNA degradation rate (min-1)
1.0
0.5
0.0
*
***
*
***
rne: WT
WT
rne3071
Temp: 30°C
43.5°C
Binding sites:
N-terminal catalytic region
MTS
RhlB, Enolase, PNPase
WT RNase E
1
529
592
1061
RNase E ΔMTS
1
529
RNase E (1-592)
1
529
592
RNase E (1-529)
1
529
0.4
0.2
0.0
ns
ns
ns
ns
ns
**
WT
ΔrhlB
Δpnp
RNase E (1-592)
RNA degradosome

### Figure S3

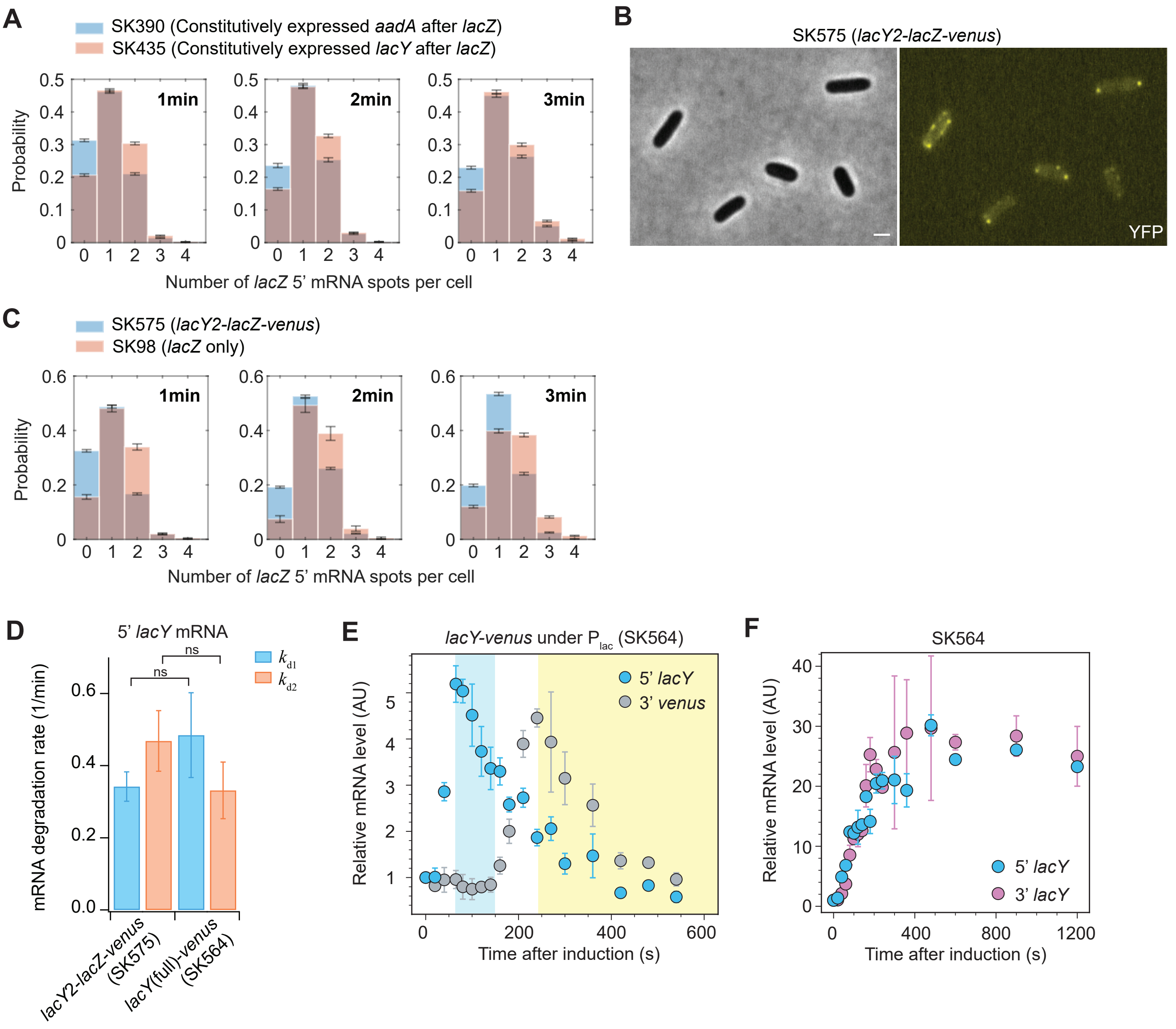

A
SK390 (Constitutively expressed aadA after lacZ)
SK435 (Constitutively expressed lacY after lacZ)
Probability
1min
2min
3min
Number of lacZ 5' mRNA spots per cell
B
SK575 (lacY2-lacZ-venus)
YFP
C
SK575 (lacY2-lacZ-venus)
SK98 (lacZ only)
Probability
1min
2min
3min
Number of lacZ 5' mRNA spots per cell
D
5' lacY mRNA
mRNA degradation rate (1/min)
ns
ns
$k_{d1}$
$k_{d2}$
lacY2-lacZ-venus (SK575)
lacY(full)-venus (SK564)
E
lacY-venus under $P_{lac}$ (SK564)
5' lacY
3' venus
Relative mRNA level (AU)
Time after induction (s)
F
SK564
5' lacY
3' lacY
Relative mRNA level (AU)
Time after induction (s)

### Figure S4

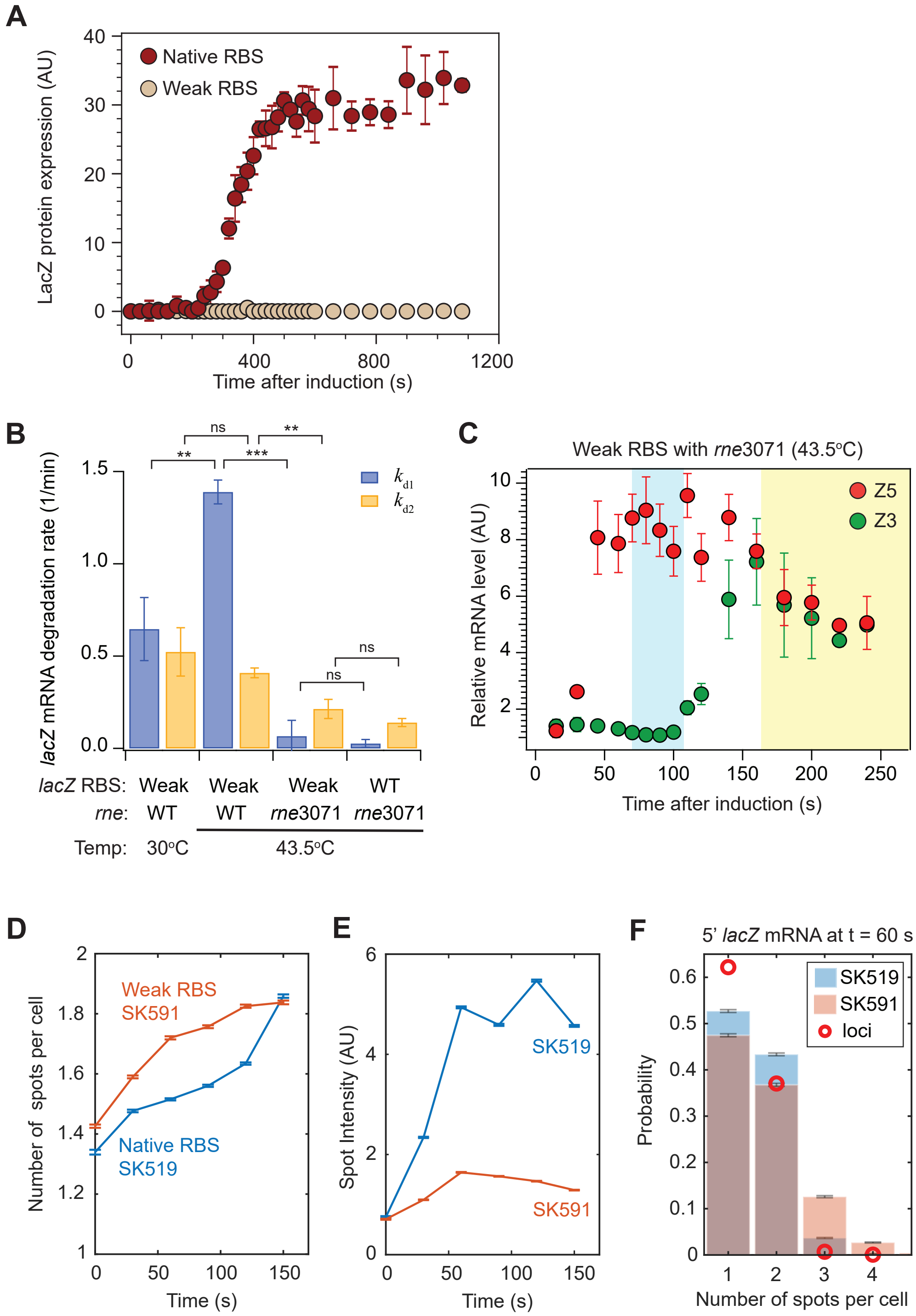
A
LacZ protein expression (AU)
Native RBS
Weak RBS
Time after induction (s)
B
lacZ mRNA degradation rate (1/min)
ns
**
***
$k_{d1}$
$k_{d2}$
lacZ RBS: Weak Weak Weak WT
rne: WT WT rne3071 rne3071
Temp: 30°C 43.5°C
C
Weak RBS with rne3071 (43.5°C)
Relative mRNA level (AU)
Z5
Z3
Time after induction (s)
D
Number of spots per cell
Weak RBS SK591
Native RBS SK519
Time (s)
E
Spot Intensity (AU)
SK519
SK591
Time (s)
F
5' lacZ mRNA at t = 60 s
Probability
SK519
SK591
loci
Number of spots per cell

### Figure S5

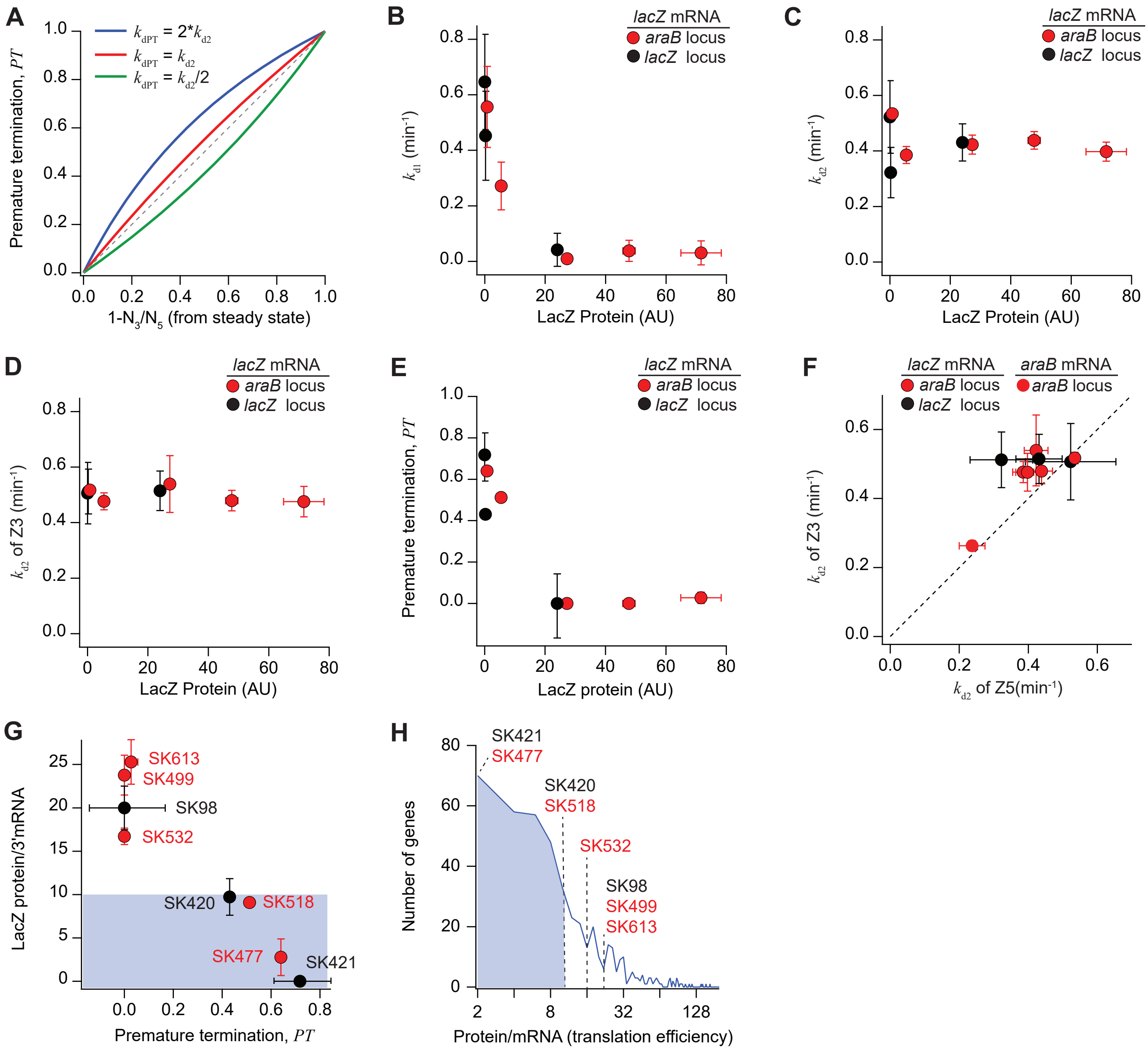

A
B
C
D
E
F
G
H
Premature termination, PT
1-N3/N5 (from steady state)
kdPT = 2*kd2
kdPT = kd2
kdPT = kd2/2
kd1 (min-1)
kd2 (min-1)
kd2 of Z3 (min-1)
kd2 of Z5(min-1)
LacZ Protein (AU)
LacZ protein (AU)
lacZ mRNA
araB mRNA
araB locus
lacZ locus
LacZ protein/3'mRNA
Premature termination, PT
SK613
SK499
SK98
SK532
SK420
SK518
SK477
SK421
Number of genes
Protein/mRNA (translation efficiency)

### Figure S6

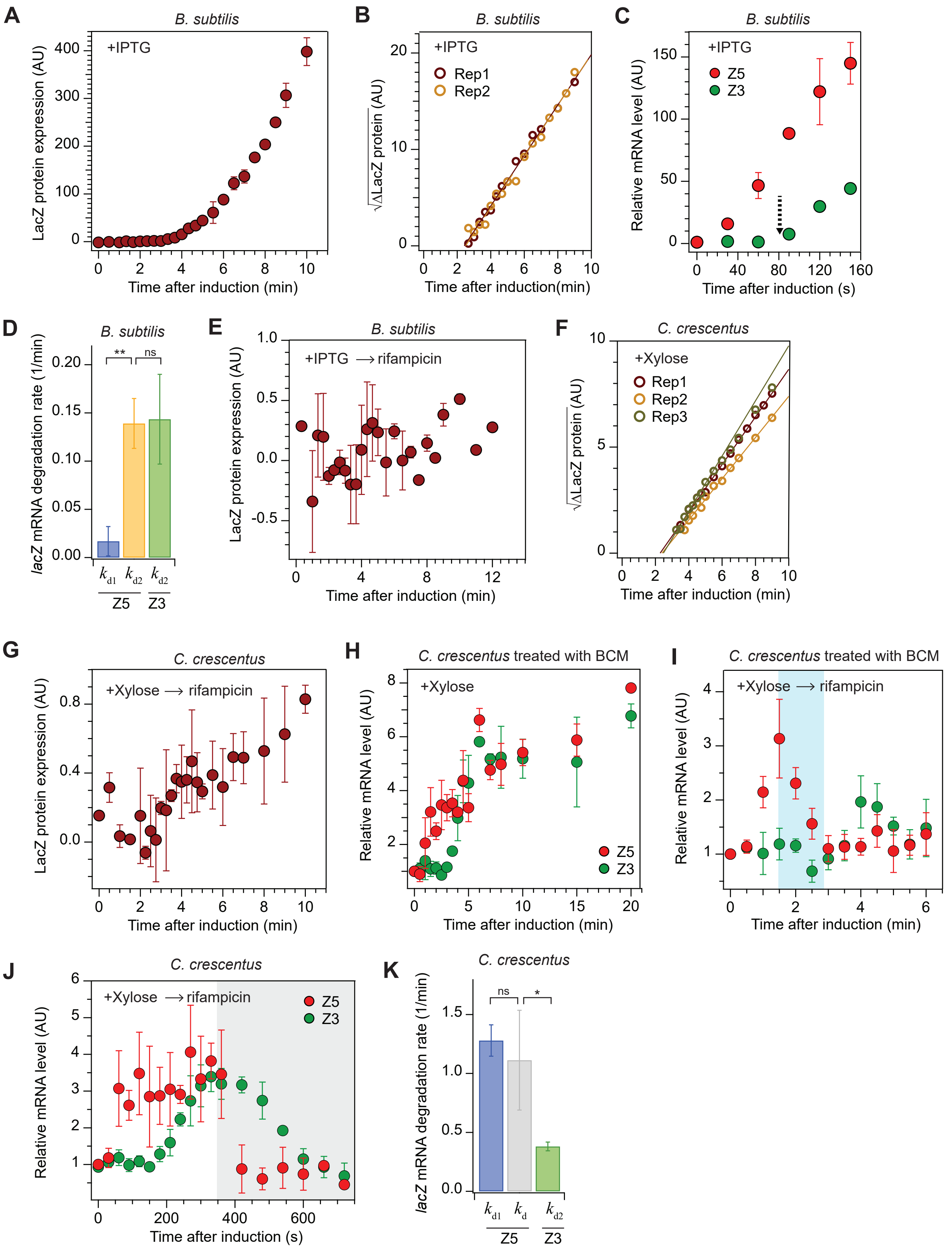

A
B. subtilis
+IPTG
LacZ protein expression (AU)
Time after induction (min)
B
B. subtilis
+IPTG
Rep1
Rep2
√ΔLacZ protein (AU)
Time after induction(min)
C
B. subtilis
+IPTG
Z5
Z3
Relative mRNA level (AU)
Time after induction (s)
D
B. subtilis
lacZ mRNA degradation rate (1/min)
**
ns
kd1
kd2
kd2
Z5
Z3
E
B. subtilis
+IPTG ⟶ rifampicin
LacZ protein expression (AU)
Time after induction (min)
F
C. crescentus
+Xylose
Rep1
Rep2
Rep3
√ΔLacZ protein (AU)
Time after induction (min)
G
C. crescentus
+Xylose ⟶ rifampicin
LacZ protein expression (AU)
Time after induction (min)
H
C. crescentus treated with BCM
+Xylose
Z5
Z3
Relative mRNA level (AU)
Time after induction (min)
I
C. crescentus treated with BCM
+Xylose ⟶ rifampicin
Relative mRNA level (AU)
Time after induction (min)
J
C. crescentus
+Xylose ⟶ rifampicin
Z5
Z3
Relative mRNA level (AU)
Time after induction (s)
K
C. crescentus
lacZ mRNA degradation rate (1/min)
ns
*
kd1
kd
kd2
Z5
Z3
